# Strong Elastic Protein Nanosheets Enable the Culture and Differentiation of Induced Pluripotent Stem Cells on Microdroplets

**DOI:** 10.1101/2023.06.22.546128

**Authors:** Elijah Mojares, Alexandra Chrysanthou, Julien E. Gautrot

## Abstract

Advances in stem cell technologies, revolutionising regenerative therapies and advanced in vitro testing, require novel cell manufacturing pipelines able to cope with scale up and parallelisation. Microdroplet technologies, which have transformed single cell sequencing and other cell-based assays, are attractive in this context, but the inherent soft mechanics of liquid-liquid interfaces is typically thought to be incompatible with the expansion of induced pluripotent stem cells (iPSCs), and their differentiation. In this work, we report the design of protein nanosheets stabilising liquid-liquid interfaces and enabling the adhesion, expansion and retention of stemness by iPSCs. We use microdroplet microfluidic chips to control the formulation of droplets with defined dimensions and size distributions and demonstrate that these sustain high expansion rates, with excellent retention of stem cell marker expression. We further demonstrate that iPSCs cultured in such conditions retain the capacity to differentiate into cardiomyocytes and demonstrate such process on droplets. This work provides clear evidence that local nanoscale mechanics, associated with interfacial viscoelasticity, provides strong cues able to regulate and maintain pluripotency, as well as to support commitment in defined differentiation conditions. Microdroplet technologies appear as attractive candidates to transform cell manufacturing pipelines, bypassing significant hurdles paused by solid substrates and microcarriers.

## Introduction

Since the early development of cell reprogramming and demonstration of somatic cell nuclear transfer [1,2], the field of induced pluripotent stem cells (iPSCs) has progressed considerably, in particular with the identification of a minimal set of transcription factors (Oct4, Sox2, cMyc, and Klf4) allowing the enhancement of the efficiency of the process [3]. Reprogramming was further refined through the application of different transfection protocols (e.g. the use of non-integrating agents such as episomal vectors and Sendai viruses (SeV) [4,5]) and the identification of alternative factors enabling effective reprogramming (e.g. replacement of cMyc and Klf4 with nanog and Lin28 [6,7]). These development allowed the identification of reprogramming strategies that are considered to be compliant with current good manufacturing practices (cGMP) and translation of iPSC technologies [8].

Reprogrammed iPSCs have the potential to differentiate into all cell types, without introducing ethical concerns associated with embryonic stem cells, making them a unique tool for the development of new technologies for regenerative medicine, drug screening, and disease modelling [9]. This includes differentiation into neuronal, endothelial, osteochondral, epidermal and cardiac lineages, amongst many examples. Several allogenic iPSC-based therapies are already undergoing clinical trials, for example targeting ophthalmic diseases, neurological disorders, cardiac repair and diabetes [10]. An autologous iPSC therapy trial was also developed in Japan, with two patients receiving transplantation of autologous iPSC-derived retinal cells to treat macular degeneration [11]. After one year, one of these patients did not display signs of complications, rejection or unexpected proliferation suggesting that such transplantation may be clinically safe and effective, although this remains to be fully established [12].

Disease modelling and therapeutic screening have also benefitted tremendously from the iPSC technologies. Whether it is through the derivation of defined cell lineages that can be incorporated into advanced in vitro models, or through the formation of organoids of brain [13–16], liver [17], prostate [18], and small intestine [19–21] tissues, amongst others, these models offer unprecedented potential to examine patient-specific disease development and test the efficacy and safety of personalised medicine. For example, pancreatic organoids derived from iPSCs were applied to the study of cancer progression [22]. iPSC-derived cerebral organoids were applied to study the impact Zika virus on neuro-epithelial development [16]. Furthermore, iPSC-derived cells from patients suffering from autism were used to create brain organoids to study the molecular mechanism of the disease, identifying the upregulation of FOX1 and subsequent overproduction of inhibitory neurons as a possible process underpinning this condition [15]. More complex models have also evolved, proposing the integration of multi-organ-chip systems, for example based on renal, liver, intestine, and neuronal organoids from patient-derived iPSCs and allowing the study of adsorption, distribution, metabolism and excretion mechanisms in an autologous patient-specific context [23].

Such strategies have unprecedented potential for the understanding of pathologies and the identification of personalised therapeutic strategies. However, the generation of iPSC lines, differentiation, handling, and culture times required can extend from several weeks to months, depending on the complexity of the models proposed and the availability of existing cell lines. Carrying out relevant pre-clinical studies on such models on sufficiently large patient-derived cohorts remains prohibitively difficult to coordinate and costly. Therefore, novel strategies and platforms allowing the scale up and parallelisation (to cope with even moderately large patient-derived cell libraries) of cell manufacturing and derivation of associated models is critically needed. Overall, the reproducibility, scalability, and safety of associated protocols remains challenging for the derivation of current complex in vitro models and organoid systems, for drug screening and regenerative medicine [24].

The scale up and automation of iPSC culture remains challenging. Some suspension platforms have been proposed, allowing the culture of iPSCs as embryoid bodies or spheroids, but these require relatively complex protocols for passaging, dissociation and recovery of the cell products, involving multiple centrifugation steps, long enzymatic treatments and passing the culture through narrow channels to help dissociation [25]. In addition, not all iPSCs can grow in suspension platforms. Microcarriers have also been applied to iPSC culture, promoting the adhesion of iPSCs and the growth of colonies [26]. These include diethyl aminoethyl-based DEAE-Sephadex that were applied to the culture of embryonic epidermal and lung epithelial cells [27,28], gelatin-based microcarriers CultiSpher-S and Cytodex-3 applied to the culture and lineage-specific differentiation of mesenchymal stem cells and pluripotent stem cells [29], or thermoresponsive polymer-based microparticles (e.g. based on poly (N-isopropyl acrylamide) and other copolymers displaying lower critical solution temperatures), for the culture of bone marrow derived mesenchymal stem cells [30,31]. However, these microcarriers require the separation of resulting cell products, in particular if implantation is targeted, and contamination with microplastics as well as high costs remain important barriers to the application of these systems.

To address issues associated with cell expansion and processing using solid substrates and microcarriers, the rational design of liquid-liquid interfaces enabling cell culture at the surface of oil droplets was recently proposed [32–36]. The self-assembly of proteins and polymers at liquid-liquid interfaces was found to result in the formation of relatively dense and stiff nanosheets that display sufficiently strong and elastic mechanical properties to resist cell-mediated deformations [36–38]. These mechanical properties can be regulated by the protein denaturation process itself [39], as well as the physico-chemistry of the proteins adsorbed [33], and the co-assembly with reactive surfactants that can confer further physical crosslinking to the resulting assemblies [32,37,40]. Macromolecular design, for example through the selection of the molecular weight of the polymers adsorbed, was also found to have a critical impact on the toughness of resulting nanosheets, through the dissipation of energy by soft polymer chain segments not pinned to the interface, in turn impacting on cell-mediated fracture of the corresponding interfaces [34]. This unique nano-mechanical environment, coupled to suitable bioactive ligands, was found to not only promote cell adhesion and spreading, but also enable the expansion of a broad range of cells and the maintenance of stem cell phenotype over prolonged culture periods [41,42]. This allowed the culture of mesenchymal stem cells at the surface of microdroplets stabilised by strong bioactive protein nanosheets, termed bioemulsions, whilst retaining marker expression and their capacity to differentiate in defined conditions. However, iPSC culture at liquid-liquid interfaces and on associated bioemulsions, and the impact that such environment may have on pluripotency and capacity to differentiate, has never been explored. The impact that the nanoscale mechanics of the cell microenvironment may play on the retention of pluripotency, at liquid-liquid interfaces, remains unknown.

Here we report the culture of iPSCs at the surface of liquid substrates. We examine the impact of protein nanosheet design and interfacial mechanical properties on the expansion of iPSCs and the formation of colonies, as well as retention of pluripotency marker expression. We then investigate whether such strategy can be applied to the design of bioemulsions for the culture of iPSCs and demonstrate the proof of concept of on-droplet induction of differentiation into cardiomyocytes. We examine the capacity of bioemulsions to support differentiation and the resulting formation of functional, contractile, microtissues.

## Materials and Methods

### Materials

Cell culture medium Essential 8 (E8) Flex Medium and recombinant human vitronectin (rhVTN-N) were purchased from Thermofisher Scientific. EZ-Lift stem cell passaging reagent, Dulbecco’s phosphate buffered saline (DPBS), trichloro(1H,1H,2H,2H-perfluorooctyl) silane, L-ascorbic acid 2-phosphate (LAA2P) and *O. sativa* derived human serum albumin (HSA) were purchased from Sigma Aldrich. CHIR99021, Wnt-C59, VEGF165, BMP4, activin-A, and bFGF were purchased from Peprotech. Gentle cell dissociation reagent and APEL(2) Medium were purchased from Stem Cell Technologies. Primary antibodies used were mouse anti-Oct 3/4 (Santa Cruz Biotechnology – SC-5279), rabbit anti-nanog (Abcam – ab109250), mouse anti-E-cadherin (Cell Signalling Technology – 14472S), rabbit anti-cardiac troponin T (Abcam – ab45932), and Mouse Anti-α-actinin (Sigma – A7811). Secondary antibodies used were goat anti-mouse IgG (H+L) highly cross-adsorbed, Alexa Fluor 488 (Thermofisher – A-11029), donkey anti-rabbit IgG (H+L) highly cross-adsorbed, Alexa Fluor 647 (Thermofisher – A-31573), and donkey anti-sheep IgG (H+L) cross-adsorbed, Alexa Fluor 555 (Thermofisher – A-21436). The proteins used were β-Lactoglobulin (βLG) (Sigma – L3908) and poly(L-Lysine) (M_w_ 30-70 kDa) (Sigma – P2636). Conjugated primary antibodies used was anti-cardiac troponin 2 (TNNT2) PE (Miltenyi Biotec – 130-120-405). Viability marker Zombie Aqua (Biolegend – 423102) was used.

### Interfacial shear rheology measurements

The quantification of interfacial shear mechanical properties was evaluated via oscillatory interfacial shear rheology using a Discovery Hydrid-Rheometer (DHR-3) from TA Instruments fitted with a Du Nouy Ring geometry and a Delrin trough with a circular channel. The Du Nouy ring has a diamond-shaped cross section, a radius of 10 mm and is made of a platinum-iridium wire of 400 μm thickness. The Delrin trough was filled with 4 mL of fluorinated oil. Using axial force monitoring, the ring was positioned at the surface of the fluorinated oil, and was then lowered by a further 200 μm to position the medial plane of the ring at the fluorinated phase interface. 4 mL of the PBS solution were then gently introduced to fully cover the fluorinated sub-phase. Time sweeps were performed at a frequency of 0.1 Hz and temperature of 25°C, with a displacement of 1.0 10^-3^ rad (strain of 1 %) to follow the self-assembly of the protein nanosheets at corresponding interfaces. In each case, the protein solution (1 mg/mL) was added after 15 min of incubation and continuous acquisition of interfacial rheology data for the naked liquid-liquid interface. After each time sweep, a frequency sweep (with displacements of 1.0 10^-3^ rad) and amplitude sweeps (at a frequency of 0.1 Hz) were carried out to examine the frequency-dependent behaviour of corresponding interfaces and to ensure that the selected displacement and frequency selected were within the linear viscoelastic region. For the evaluation of the impact of sulfo-SMCC coupling, one hour after the formation of the protein layer the aqueous phase was exchanged using a high resolution flow control system (Elveflow), without disrupting the formed nanosheet in the rheology trough. To ensure that the aqueous phase had been fully exchanged, flow was continued for 30 minutes. Sulfo-SMCC was then introduced into the aqueous phase at final concentration of 2 mg/ mL, 30 min after end of the washing period and acquisition of interfacial rheology data was continued for 1 hour. Another 30 min washing was carried out, followed by frequency sweep (with displacements of 1.0 10^-3^ rad), stress relaxation experiment (0.5 % strain) and amplitude sweeps (at a frequency of 0.1 Hz).

### Induced Pluripotent Stem Cell Culture

The HPSI0114i_vabj3 iPSC cell line was obtained from the HipSci iPSC cell bank. These cells were cultured with E8 Flex Medium rhVTN-N coated plates. Plates or substrates were incubated with rhVTN-N (10 μg/mL) for at least 1 hour before seeding of cells. For passaging, iPSCs were first washed with DPBS. iPSCs were detached from well plates using the EZ-Lift stem cell passaging reagent, as recommended by the supplier. Briefly, cells were incubated with EZ-Lift on an orbital shaker at 37°C, in a 5% CO_2_ incubator. After 4 minutes, plates were tapped rapidly until clumps of cells were visibly dislodged. The cell suspension was then transferred to a centrifuge tube with medium. The resulting cell suspension was then centrifuged at 120 × g for 3 minutes. The supernatant was aspirated and the cell pellet was resuspended in E8 Flex medium.

For seeding iPSCs on interfaces as single cells, plates were first pretreated with E8 flex medium supplemented with 10 μM Y-27632 for 1 hour. The concentration of ROCK inhibitor, Y-27632, is routinely used for iPSC culture and is recommended by HipSci. The plates were then washed once with DPBS and incubated for 8 minutes at 37°C with gentle cell dissociation reagent. With a 5 mL stripette, the cells were pipetted up and down at maximum speed 3 times. The resulting cell suspension was further diluted with equal amount of medium and centrifuged at 120 × g for 3 minutes. The supernatant was aspirated and the cell pellet was resuspended with 1 mL medium with a P1000 micropipette to ensure the cells were single cells. The resulting cell suspension was further diluted and counted.

### EB Formation

Polyvinyl alcohol (PVA) was dissolved in E8 flex medium overnight to achieve a concentration of 4% PVA. This was then filtered and supplemented with 10 μM Y-27632. iPSCs were dissociated with Gentle Cell Dissociation Reagent according to manufacturer’s instructions. After centrifugation, cells were counted. An aliquot of the cell suspension was then taken and centrifuged at 120 × g for 3 minutes. The cell pellet was then resuspended in E8 flex with PVA and Y-27632 to achieve a cell concentration of 10^6^ cells/mL. A petri dish was then filled with sterile water. The lid was then placed upside down and multiple 10 μL drops of the cell suspension was then placed on the lid. The lid was then placed back on the petri dish resulting in hanging droplets. After 24 hours, 30 embryoid bodies were transferred to a well of an ultra-low attachment 6-well plate filled with E8 flex medium.

### Generation of interfaces

Pre-treatment of the wells to generate flat interfaces was needed following protocols adapted from the literature [38]. Briefly, an M24 well plate was first plasma treated for 10 minutes (Henniker Plasma HPT-200). Two solutions were prepared. For solution A, per well to be treated, 300 μL of ethanol or methanol was mixed with 12 μL of triethylamine. For solution B, 3.6 mL of ethanol or methanol was mixed with 144 μL of trichloro(1H,1H,2H,2H-perfluorooctyl) silane. After plasma treatment, Solution A was added and shaken lightly to evenly coat the wells. Solution B was then added and immediately sealed with parafilm. The reaction was incubated overnight at room temperature.

To prepare the flat interfaces, the reaction mixture was aspirated and wells were briefly washed with 70% ethanol. The wells were then washed with DPBS three times and left to dry briefly. Novec oil was added at 500 μL per well and 2 mL of DPBS was added to each well. For PLL interfaces, DPBS pH 10.5 was used instead of pH 7.4. This was then incubated at the CO2 incubator for 20 minutes. Any air bubbles were removed and concentrated protein solutions were added. The stock solution for βLG and poly(L-lysine) was 100 mg/mL and 10 mg/mL respectively. The working concentration for these two proteins were 1 mg/mL and 100 μg/mL respectively. After 1 hour, the excess protein was washed off with DPBS 6 times, leaving behind an estimated 500 μL of aqueous solution. If sulfo-SMCC was to be added, 500 μL of 2 mg/mL sulfo-SMCC was added to the resulting interface. Sulfo-SMCC was then washed off 6 times with DPBS after 1 hour. The interfaces were then incubated with rhVTN-N (10 μg/mL) for 1 hour and washed off 3 times with E8 flex medium. Per well, 500 μL of E8 flex medium supplemented with 20 μM Y-27632 were added – resulting in an aqueous solution with E8 Flex medium supplemented with 10 μM Y-27632. Cells were dislodged from their plates, resuspended in E8 flex medium with Y-27632, counted with a haemocytometer, and plated at 40,000 cells per well.

For pinned droplets, glass slides were plasma treated for 10 min (Henniker Plasma HPT-200). The plasma treated glass slides were then incubated for 1 h with 5% (v/v) Trichloro(1H,1H,2H,2H-perfluorooctyl) silane in anhydrous ethanol or methanol. Excess silane was washed off with anhydrous ethanol or methanol and dried with nitrogen gas. For 35 mm petri dishes, 13 mm holes were cut and the fluorinated glass were stuck onto the bottom of the Petri dish with Sylgard 184 PDMS where the monomer and curing agent were mixed together at a 10:1 ratio. This was then cured for 2 hours or overnight at 70°C. Alternatively, fluorinated glass slides were attached to ibidi chambers with sticky wells.

To prepare the pinned droplets, the vessels were first washed briefly with 70% ethanol and washed 3 times with DPBS. For the petri dishes, 3 mL of DPBS was added and 10 μL of Novec 7500 oil was spread throughout the glass. For Ibidi chambers, 500 μL of DPBS or DPBS pH 10.5 were added and 7 μL of Novec 7500 oil were spread at the bottom. These were then incubated at 37°C for 20 min and air bubbles were dislodged. The concentrated protein solution was then added, at working concentrations used for flat interfaces. Washes and incubation, as for flat interfaces, were carried out for pinned droplets. The seeding density used was 13333 cells per cm^2^.

After 24 h, Y-27632 was washed off with DPBS once and E8 Flex medium three times. For pinned droplets, cells were then grown for 4 days before being fixed and immunostained. For flat interfaces, immediately after removing the excess Y-27632, the cells were live imaged for 48 h (Lumascope 720).

### Microfluidic Device Fabrication

The microfluidic device was fabricated using typical soft lithography techniques. Briefly, a silicon wafer is cleaned with acetone and isopropanol. The silicon wafer was then dried with nitrogen gas. SU-8 2050 /SU-8 2100 (100 µm vs 300 µm height) is then spin coated onto the silicon wafer. It was then soft baked on a levelled hotplate. After cooling down, a glass plate with the photomask taped onto it is placed on top of the silicon wafer with the spin coated SU-8. The SU-8 was then exposed to UV light. Afterwards, the wafer was baked on a levelled hotplate. The wafer was then immersed in a dish containing the developer solution. After some time, the wafer was rinsed with isopropanol. This was repeated until no white deposits was visible. The wafer was then dried under nitrogen gas and placed in a Petri dish. Depending on the desired thickness of the SU-8 masters, the spin speed and pre-exposure bake times were adjusted. For 100 µm thick masters, a spin speed of 1500 rpm was used and the pre-exposure bake times were 65°C for 5 min and 95°C for 10 min. For thicker moulds of around 300 µm, a spin speed of 1000 rpm was used and pre-exposure bake times were 65°C for 7 min and 95°C for 40 min.

PDMS monomer and curing agent were mixed together at a 10:1 ratio. This was then mixed together until opaque. This was then degassed several times in a desiccator and placed under vacuum until the mixture was clear and transparent. The mixture was then poured onto the petri dish containing the wafer. This was then baked at 80°C for 2 h. The hardened PDMS was then cut with a scalpel. A biopsy punch (1.5 mm) was used to bore the inlets and outlets on the device. Dust was removed from the device with tape. The device and glass slides were further cleaned with water and isopropanol and dried with nitrogen gas. The device and glass slide were then treated with plasma for 2 min at 100% power. The device and glass slide were then pressed onto each other to bond them together.

To assemble the device together, PTFE tubing (1.0 mm OD) Tygon Microbore tubing (0.020” × 0.060” ID × OD) (Cole-Parmer) were inserted directly into the inlets and outlets of the PDMS chip. These were stuck in place with a resin (Bostik Evo-Stik Ultra Strong Control Adhesive).

### Microdroplet Generation with Microfluidic Devices

The device was initially sterilized by flowing 70% ethanol through the device. An OB1 MKIII+ pressure controller from Elveflow was used to regulate the pressure within channels (in the range of 0 to 2000 mbar). Centrifuge tubes were connected to each pressure channel via a metal adaptor with 1/4”-28 UNF(F) thread. A barbed fitting was attached to the metal cap where a 3/32” ID tygon tubing can then be connected to the cap and the pressure channel. A filter was placed at the other end of the tubing before connecting it to the OB1 MKIII+. The tubings of the devices were then connected to the reservoirs via 1/4”-28 UNF(M) flat bottom flangeless fittings. The pressure was then controlled using the Elveflow Software Interface (ESI).

After sterilization, the protein solution were flowed through the aqueous channel and air filters through the central channel for 20 min. βLG (1 mg/mL) solution in PBS pH 7.4 was flowed through the aqueous channel while the Novec 7500 oil was flowed through the central channel. The resulting microdroplets were then collected in a reservoir containing the βLG solution and incubated for an hour. After incubation for an hour, the microdroplets were washed with PBS pH 7.4 six times to remove excess PLL and to return the pH to the physiological pH.

The microdroplets were then coated with Sulfo-SMCC (2 mg/mL) and incubated for an hour. The microdroplets were then washed six times with PBS pH 7.4. The microdroplets were then coated with vitronectin (10 µg/mL) and incubated for another hour. After an hour, the microdroplets were washed with PBS three times and resuspended in media. Resulting microdroplets were imaged with an inverted microscope.

### Cardiomyocyte Differentiation

iPSCs were seeded on microdroplets and tissue culture plastic controls in E8 flex medium supplemented with Y-27632, a ROCK inhibitor. After 24 h, the medium was replaced with E8 flex medium. After 4 days of culture, unless otherwise stated, cardiomyocyte differentiation was started. The cardiomyocyte differentiation protocol was adapted from the literature [43]. Briefly, a chemically defined medium composed of 3 reagents (CDM3) was used as the basal medium. It is composed of RPMI 1640 medium (w/ glucose), O. sativa derived human serum albumin (HSA), and L-ascorbic acid 2-phosphate (LAA2P). To initiate mesoderm differentiation, the cells grown on monolayers, either on microdroplets or TCP, were incubated in CDM3 supplemented with CHIR99021 for 2 days. Afterwards, the Wnt pathway was inhibited with 5 μM Wnt-C59. After 2 days, the medium was refreshed with CDM3 only and changed every other day. On the 10th day, metabolic selection was performed with CDM3L medium wherein RPMI 1640 without D-glucose was used and instead is supplemented with 5 mM sodium D-lactate. The medium was replaced with CDM3L every other day. After 6 days of metabolic selection, cultures were recovered for analysis.

### Immunostaining and Microscopy

The media was first aspirated, and cells were washed with PBS. The cells were then fixed with 4% PFA for 10 min. Permeabilization was performed with 0.2% Triton-X100. Blocking was performed with 3% bovine serum albumin for 1 h. The primary antibody was diluted in 3% BSA solution then added to the cells and incubated overnight at 4°C. The cells were then washed three times with PBS. The secondary antibody, along with Alexa Fluor-555 phalloidin and DAPI, was diluted in 3% BSA then added to the cells and incubated for another hour. The cells were then washed with PBS three times. For cells on cover slips, mounting was carried out on glass slides with Fluoromount-G (SouthernBiotech). For cells on emulsions, cells were transferred to an 8-well µ-Slide containing PBS solution. Cells on pinned droplets were imaged directly through the substrate, using a Zeiss LSM710 confocal microscope. For each pinned droplet setup, 8 images were taken.

### Live Imaging and Image Analysis

Live imaging was performed with a Lumascope 720 set up in a CO_2_ incubator. Cells were incubated at 37°C and 5% CO_2_ and imaged for 48 h every hour. Per well, 3 regions of interest (ROIs) were selected and 2×2 tiles were imaged. Three wells were made for each interface of interest, which brings the total ROIs to 9. After 48 h of imaging, the tiles for each ROI were then stitched together. The colonies formed and after 48 h were manually selected and the areas were measured. The stitched tiles should at least be covered with cells by 30%. Selected ROIs with less than 30% were excluded from the analysis.

### Image Analysis of Pluripotency Markers

For image analysis, maximum projections were made from the DNA, Oct3/4, and Nanog channels. Using ImageJ, these images were then each individually thresholded. The thresholded maximum projection of the DNA’s particles were analyzed and was set as the ROIs. These ROIs were superimposed onto the maximum projections of the Oct3/4 and Nanog channels’ maximum projections to measure the mean fluorescence for each ROI. These were then plotted as a histogram for each setup and the nuclei with a mean fluorescence intensity above 160 a.u. were counted as positive cells.

### Flow cytometry

Cells cultured on droplets or as embryoid bodies were transferred to a 15 mL centrifuge tube and washed once with PBS. Cells were then dissociated using Accutase for 5 minutes with gentle shaking on an orbital shaker. Accutase was then neutralised with E8 flex medium and centrifuged at 120 × g for 3 min. The supernatant was aspirated and the cells were resuspended in 1 mL of cold FACS buffer (10% FBS and 0.1% sodium azide in DPBS). The cell suspension was transferred to a new tube and centrifuged at 120 × g for 3 min. Cells were then resuspended in 100 µL of cold FACS buffer and transferred to a V-bottom plate. The plate was then centrifuged at 300 × g for 5 min at 4°C. Antibodies for surface markers were diluted in FACS buffer. Surface marker staining was carried out by incubating the cells in the antibody cocktail for 30 min. After incubation, cells were washed twice with PBS and incubated in Zombie Aqua (1:500) in PBS for 15 min. Cells were then washed with the FACS buffer. Fixing and permeabilization was performed using True-Nuclear Transcription Buffer Set (Biolegend) and according to manufacturer’s protocol. For intracellular staining, antibodies were diluted in the permeabilization buffer. Cells were incubated for 30 min at room temperature in the dark with the intracellular antibody cocktail. After incubation, cells were washed twice with FACS buffer.

### Statistical Analysi

Unless otherwise specified, statistical analysis was performed using one-way ANOVA with Tukey post hoc analysis. n.s., not significant; *, p < 0.05; **, p < 0.01; ***, p < 0.001.

## Results and Discussion

### Engineering of protein nanosheet-stabilised bioemulsions for the culture of iPSCs

In order to design bioemulsions that would support the culture of iPSCs and their differentiation, we explored a range of different liquid-liquid interfaces stabilised by two different types of protein nanosheets (Figure 1). The first was based on poly(L-lysine) (PLL), co-assembling with the co-surfactant pentafluoro benzoyl chloride (PFBC) [34,42]. Indeed, this assembly allows the subsequent adsorption of moderately acidic proteins such as fibronectin (pI 5.8-6.3 [44]) and vitronectin (pI 4.7-5.3 [45]) at neutral pH, without further bioconjugation step. The second was based on the globular protein β-lactoglobulin (βLG), which was previously found to form relatively elastic nanosheets even in the absence of co-surfactants [39]. The introduction of the co-surfactant PFBC in the formulation of βLG nanosheets, and their bioconjugation with the hetero-bifunctional sulfo-SMCC were also proposed to further modulate interfacial mechanical properties and enable biofunctionalisation with vitronectin, respectively.

**Figure 1.**
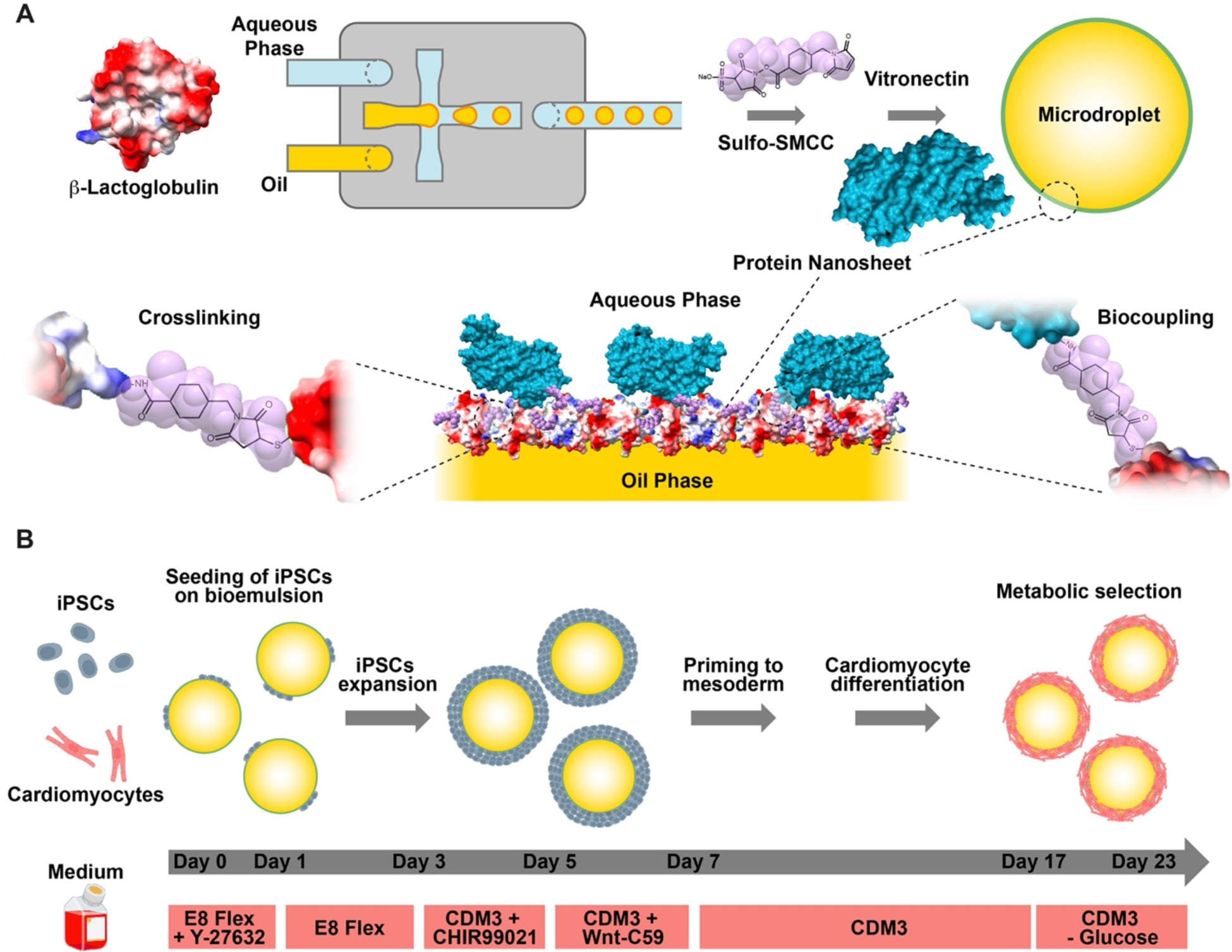
Engineering of protein nanosheet-stabilised bioemulsions for the culture of iPSCs. A. Schematic representation of the production of well-defined protein nanosheet-stabilised microdroplets generated using a flow-focusing microfluidic device, followed by their crosslinking and biofunctionalisation with vitronectin. B. Methodology applied for the expansion of iPSCs and their differentiation into cardiomyocytes, on nanosheet-stabilised bioemulsions.

The self-assembly and interfacial mechanics of protein nanosheets at the interface between the fluorinated oil Novec 7500 and phosphate buffer saline solutions was investigated via interfacial shear rheology (Figure 2) [46,47]. Using a du Noüy ring to probe interfacial mechanical properties is particularly sensitive to in plane mechanics of liquid-liquid interfaces as this can rely on very sensitive rheology-based measurements and is relatively insensitive to surface tension, as negligeable changes in surface area are associated with the in-plane deformation of interfaces. In contrast, force probe microscopy was found to be more sensitive on changes in surface areas and the disjoining pressure resulting from the probe indentation [48]. Protein solutions were injected in the aqueous phase, after equilibration of the pristine liquid-liquid interfaces and quantification of background drag levels, to monitor the kinetics of assembly (Figure 2B). Initial adsorption was found to be taking place withing a few seconds of injection, with interfacial shear moduli rapidly raising in all cases over the next 100-200 s to levels of 10-20 mN/m. In the case of βLG nannosheet assembly, this was followed by the establishment of a plateau over the next 1000 s, whereas in the case of PLL nanosheets, this was associated with a slower second step kinetics and a plateau establishing within the next 5000 s. This difference likely stems from the previously proposed mechanism of PLL nanosheet assembly, resulting from the coupling of PFBC hydrophobic residues to PLL molecules at interfaces, followed by further coupling and slower maturation of hydrophobic rigid domains able to form extended networks over large mm to cm scales [32,34].

**Figure 2.**
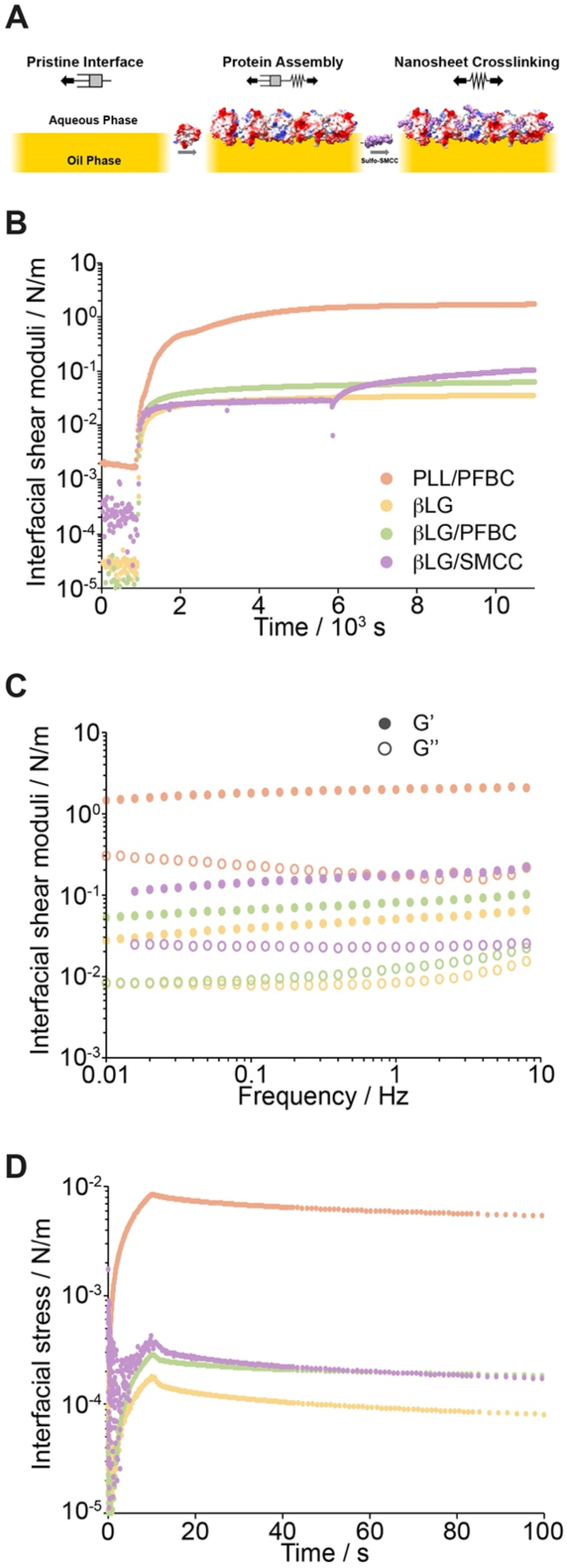
Interfacial shear rheological properties of protein nanosheet-stabilised liquid-liquid interfaces. A. Schematics of the sequential βLG adsorption and sulfo-SMCC crosslinking. B. Evolution of the interfacial storage shear modulus during the assembly of protein nanosheet at the interface between Novec 7500 oil and PBS (frequency, 1Hz; strain 10^-3^ rad). Protein injection (at 900 s; 1 mg/mL for βLG and 100 μg/mL for PLL) is indicated with a black arrow. For PLL/PFBC and βLG/PFBC, the concentration of PFBC supplemented to the oil phase was 10 μg/mL. In the case of βLG/SMCC, following exchange of the aqueous phase to remove free βLG (indicated by black arrow), SMCC was injected at a concentration of 2 mg/mL (black arrow). C. Corresponding frequency sweeps (strain 10^-3^ rad). D. Stress relaxation profiles (strain 10^−3^ rad).

Unexpectedly, sulfo-SMCC coupling was found to have a significant impact on the interfacial mechanics of βLG nanosheets, with a clear increase in interfacial shear moduli observed upon injection of sulfo-SMCC solutions (indicated by arrows, Figure 2B). This occurred despite the washing of excess of free βLG molecules, which may have resulted in multi-layer formation and aggregation of further proteins from solution. Therefore, it is proposed that sulfo-SMCC leads to further crosslinking of βLG nanosheets, either through direct tethering (maleimide moieties may couple to free cysteines and lysine residues), or via hydrophobic forces associated with its relatively apolar cyclohexyl groups.

Frequency sweeps confirmed resulting changes in interfacial shear mechanics, with broad viscoelastic regions where interfacial shear moduli were found to be relatively independent from deformation frequencies (Figure 2C). In line with the evolution of interfacial shear moduli observed from time sweeps, the interfacial storage shear modulus of PLL-stabilised interfaces was found to be considerably higher than that of βLG nanosheets, near 2 N/m. βLG assemblies alone displayed interfacial shear storage moduli in the range of 30-50 mN/m, and this increased to 50-70 mN/m, when assembly was allowed in the presence of PFBC. When βLG assembly was followed by sulfo-SMCC coupling, interfacial shear storage moduli rose further to 100-200 mN/m. In all cases, interfacial loss moduli were found to be approximately 1 order of magnitude lower than the corresponding storage components, indicating relatively strong elastic behaviours. This was also in agreement with stress relaxation profiles, which showed modest relaxation over the time scales investigated (Figure 2D). Therefore protein nanosheet properties with a relatively broad range of mechanical properties were generated, for further evaluation of iPSC adhesion and proliferation.

The formation of homogenously dispersed microdroplets stabilised by protein nanosheets and their biofunctionalisation with vitronectin was investigated next (Figure 3). Vitronectin was selected, as it is typically found to promote rapid iPSC adhesion and proliferation at solid or hydrogel interfaces [8,26]. To generate microdroplets with defined and reproducible diameters, a flow focusing microfluidic device was selected (Figure 1A and Supplementary Figure S1). This device was found to result in excellent control of microdroplet production with fluorinated, polymeric and lipid surfactants [49–52]. Similarly, βLG was found to allow excellent control of microdroplets generated with such flow focusing system, depending on the pressure and flow rate applied to the aqueous phase, with diameters ranging from 125 to 235 μm (Figure 3A-B). The lower droplet diameter achieved at the lowest aqueous phase pressure of 700 mbar may reflect a change in microdroplet formation regime [53]. In turn, treatment of the resulting droplets with sulfo-SMCC, followed by incubation with vitronectin and immuno-staining evidenced clear assembly at associated oil-water interfaces and biofunctionalisation of nanosheet-stabilised microdroplets (Figure 3C).

**Figure 3.**
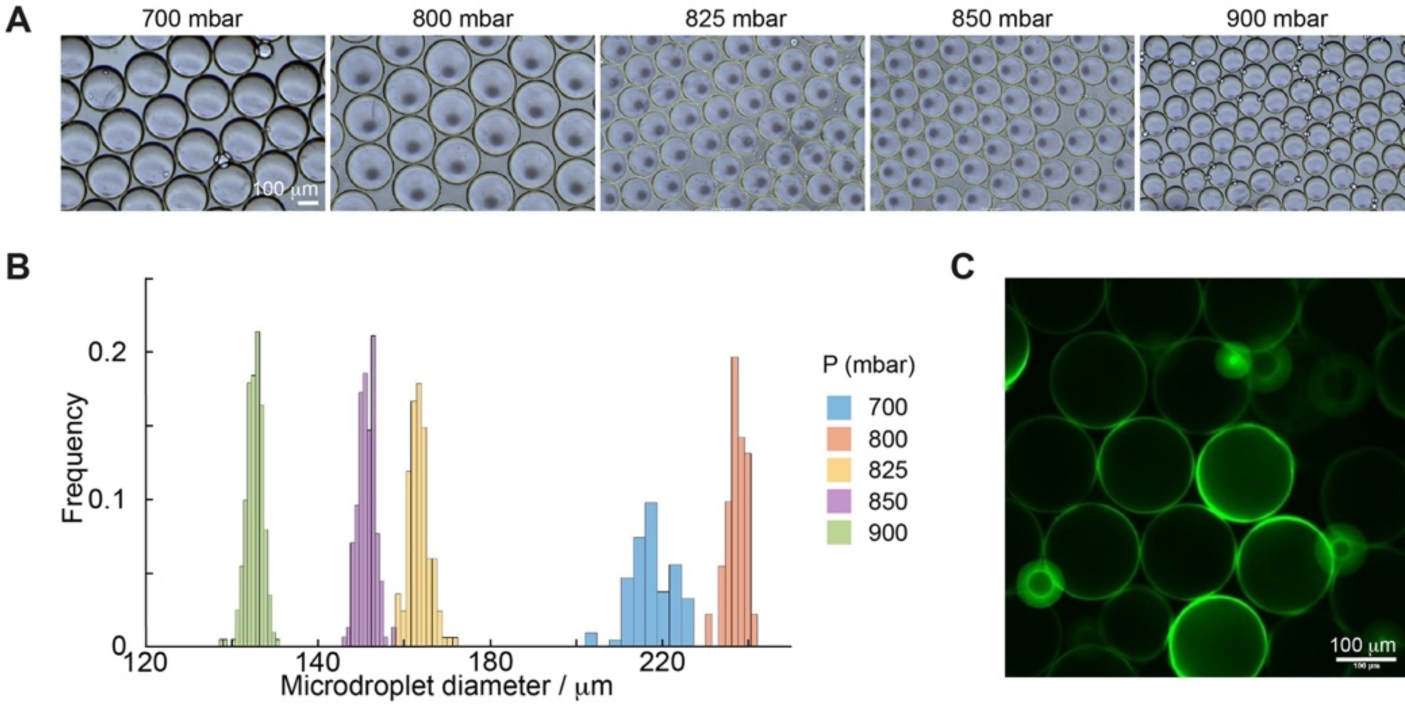
Generation of biofunctionalised nanosheet-stabilised microdroplets. A. Images of βLG-stabilised microdroplets (Novec 7500 in PBS) generated using a flow-focusing microfluidic chip, at different aqueous flow rates (controlled via the associated pressure, reported in mbar; βLG concentration of 1 mg/mL was used). B. Corresponding quantification of resulting size distributions. C. Fluorescence microscopy image of βLG-stabilised microdroplets functionalised with sulfo-SMCC (2 mg/mL) and vitronectin (10 μg/mL).

### Maintenance of the pluripotency of iPSCs upon culture at protein nanosheet-stabilised liquid-liquid interfaces

To assess early cell adhesion, colony formation and expansion at the surface of liquids, iPSCs were first seeded on flat Novec 7500 oil interfaces and droplets pinned at the surface of hydrophobized glass coverslips (to allow imaging). This allowed presenting relatively flat liquid-liquid interfaces to image colony expansion. Upon initial adhesion, iPSCs rapidly expanded at protein nanosheet-stabilised liquid-liquid interfaces (Figure 4A). After 48h of culture, large colonies could be seen at oil interfaces stabilised by βLG nanosheets functionalised by sulfo-SMCC, with similar morphologies and comparable sizes to colonies formed on vitronectin-coated glass substrates (Figure 4A/B). In contrast, βLG nanosheets alone and βLG nanosheets crosslinked by PFBC prior to functionalisation with sulfo-SMCC did not support as extensive iPSC colony formation. Similarly, PLL nanosheets crosslinked with PFBC did not support the formation of large iPSC colonies. The size of iPSC colonies correlated with the initial size of adhered iPSC aggregates, 2 h after seeding (Figure 4A), with particularly poor cell adhesion and spreading on unfunctionalized and uncrosslinked βLG nanosheets. Although colonies were found to expand on all interfaces (Figure 4C), this initial poor adhesion was sufficient to prevent extensive colony formation. Similarly both βLG and PLL nanosheets crosslinked with PFBC and functionalised with vitronectin did not support as extensive initial adhesion and subsequent colony formation. This was thought to result from a level of cytotoxicity of PFBC on iPSCs, although at the concentration used (312 ng/mL), this molecule had not be found to display toxicity on primary keratinocytes, mesenchymal stem cells or HaCaT cells [38,40,42]. Therefore, these data suggest that even moderately stiff protein nanosheets with interfacial shear moduli, in the range of a few tens of mN/m display sufficiently strong mechanical properties to enable iPSC colony adhesion and proliferation, however the differential regulation of initial colony seeding impacts on their expansion at corresponding liquid-liquid interfaces.

**Figure 4.**
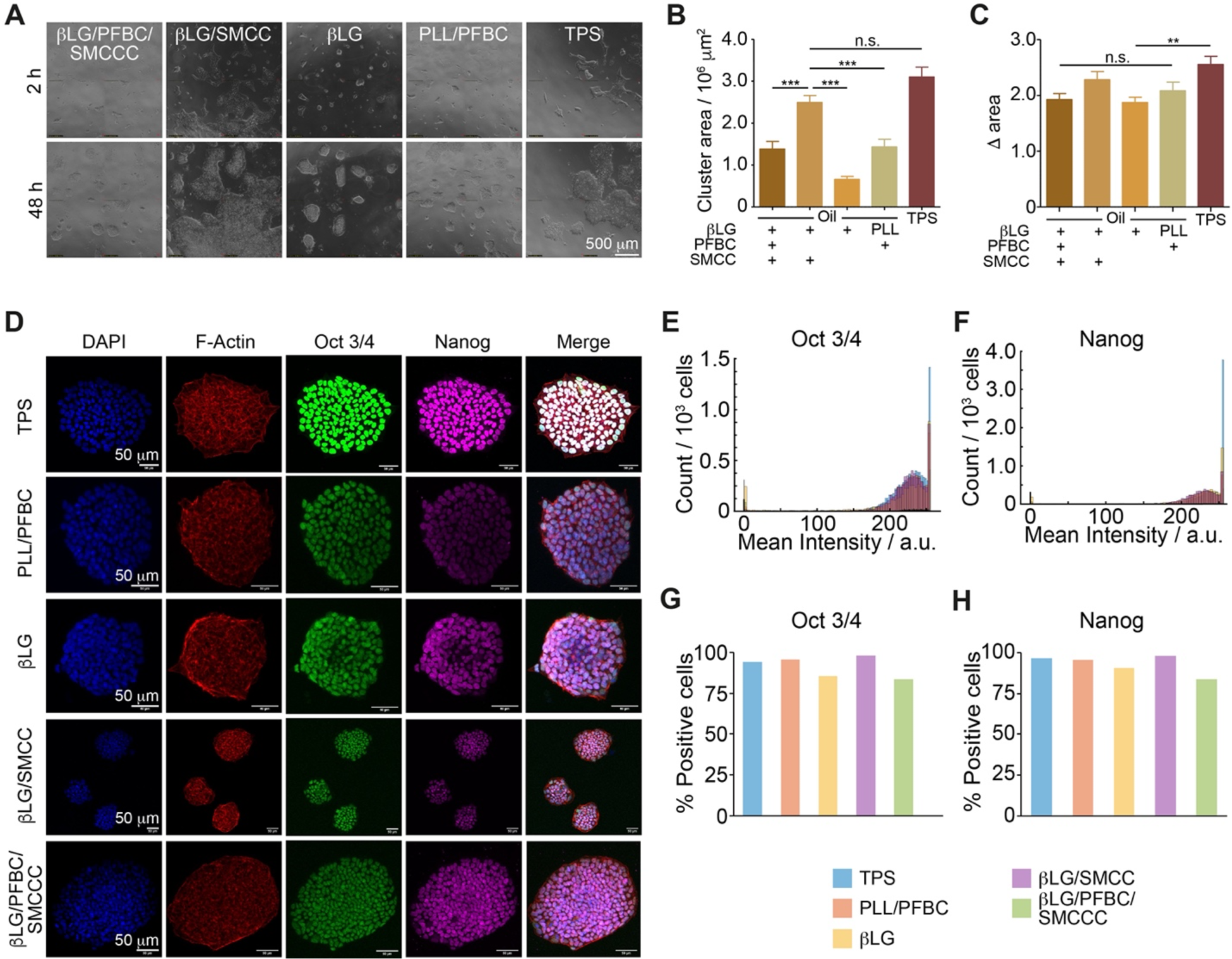
Expansion of iPSCs at nanosheet-strengthened liquid-liquid interfaces. A. Images of iPSC colonies growing at liquid-liquid interfaces and on tissue culture polystyrene (TPS) at 2 and 48 h. B and C. Corresponding quantification of changes in colony areas. 5 images taken per condition/time point; n =3. ***, p < 0.001; **, p<0.01. D. Confocal microscopy images of iPSCs growing at corresponding interfaces. Blue, DAPI; Red, Phalloidin/F-Actin; Green, Oct 3/4’ Magenta, Nanog. E-H. Corresponding quantification of mean nuclear intensities and populations of cells positive for pluripotency markers.

The maintenance of expression of the pluripotency markers Nanog and Oct 3/4 was examined next (Figure 4D-H). Immunostaining clearly confirmed that iPSC colonies growing on glass substrates or the various liquid-liquid interfaces stabilised by protein nanosheets retained high expression levels of these two transcription factors. A slight reduction in expression levels was observed for uncrosslinked and unfunctionalised βLG nanosheets and those both crosslinked and activated for biofunctionalisation, perhaps due to their lack of coupling of vitronectin and presence of PFBC, respectively. However iPSCs cultured on PLL nanosheets displayed comparable Nanog and Oct 3/4 expression to colonies growing on functionalised glass or sulfo-SMCC activated βLG nanosheets. We noted that the expression profiles of nuclei positive for these transcription factors were comparable for all interfaces (Figure 4E/F).

### Expansion and maintenance of pluripotency of iPSCs grown on selected microdroplets

Based on the excellent proliferation profiles observed for iPSCs cultured on SMCC-activated βLG nanosheets, βLG-SMCC was selected for the stabilisation of microdroplets and further evaluation of iPSC culture on bioemulsions. Droplet sizes of 150 and 230 μm were selected, based on the range achieved using flow focusing microfluidic chips (Figure 3) and in line with typical dimensions of microcarriers used for cell culture [54–56]. After functionalisation with vitronectin, iPSCs were seeded on the resulting droplet and cultured in expansion medium for 7 days. Growth was compared for droplets left to incubate in static conditions, or under dynamic flow, using an orbital shaker (70 rpm). The resulting cultures were immunostained to investigate pluripotent marker expression (Figure 5A) and iPSC proliferation was characterised by DNA quantification (Figure 5B). Initial seeding densities (evaluated at day 1) were found to be comparable on droplets to that observed on tissue culture polystyrene (TPS; Figures 5B and D). Similarly, at days 2, 4 and 7, cell counts were found to be comparable in all dynamic conditions, indicating extensive proliferation at the surface of oil droplets, whether in static or dynamic conditions (Figures 5B and C). In static conditions, cell counts on 150 μm droplets were found to be slightly lower at day 7. This effect could be associated with the increased aggregation of multiple droplets that was particularly pronounced in this condition.

**Figure 5.**
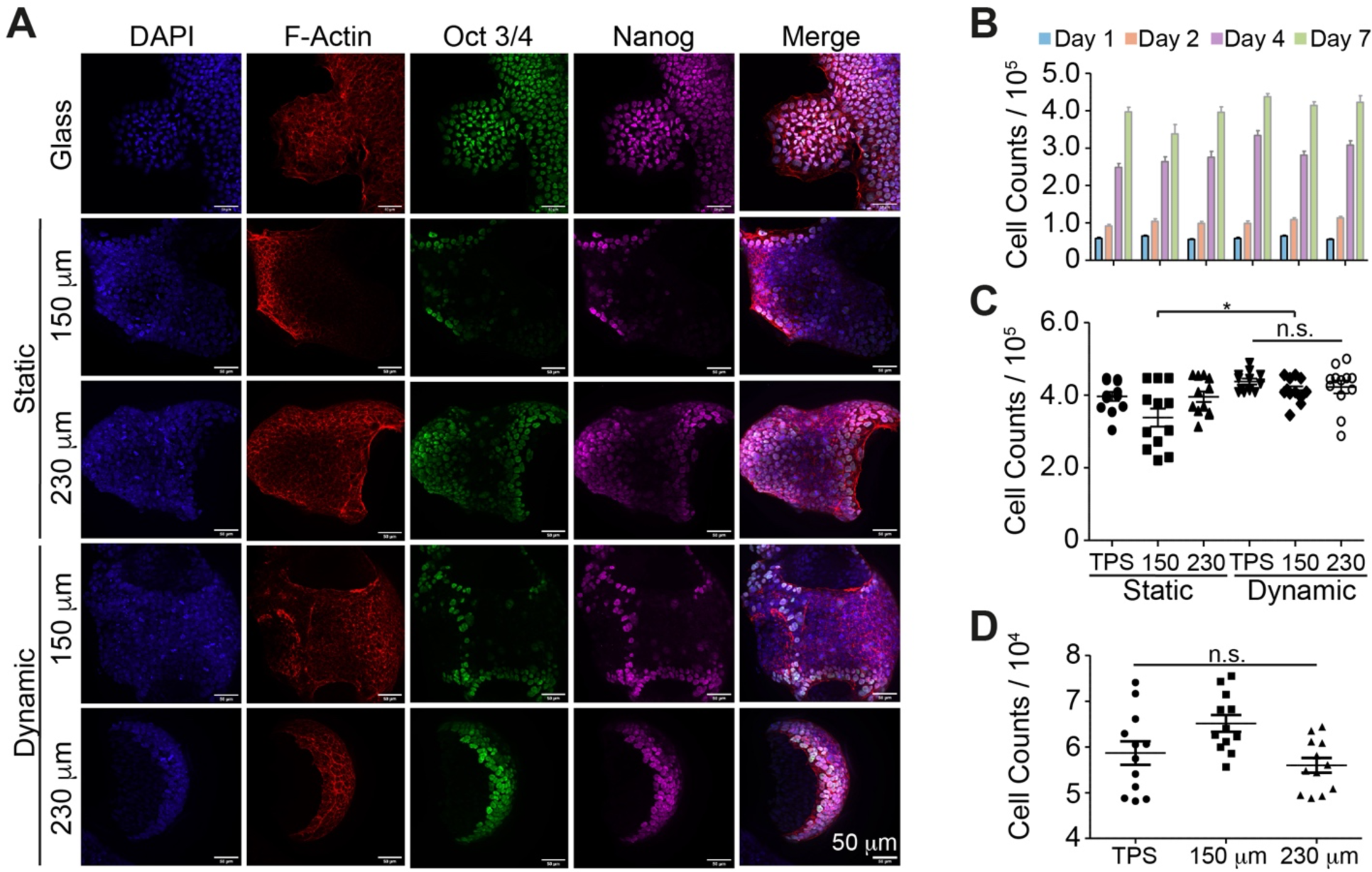
Impact of engineered protein nanosheet-stabilised bioemulsions on iPSC expansion and phenotype. A. Confocal micrographs of iPSC colonies growing on βLG-SMCC-stabilised microdroplets (diameters of 150 and 230 μm), in static and dynamic conditions. Staining for pluripotency markers Nanog (magenta) and Oct 3/4 (green). Blue, DAPI; Red, Phalloidin/F-actin. B. Proliferation profile of iPSCs grown in static or dynamic condition on TCP, 150 μm, and 230 μm βLG-SMCC microdroplets. C. Total number of cells quantified after 7 days of culture in corresponding conditions. D. Seeding efficiency of iPSCs on TCP, 150 and 230 μm βLG-SMCC microdroplets.

In addition, microscopy images of immunostained colonies growing around droplets confirmed the retention of pluripotency markers Oct 3/4 and Nanog (Figure 5A). Nuclei of iPSCs growing on bioemulsions, whether in static or dynamic conditions displayed clear localisation of these transcription factors. However, although cells remained adhered to droplets, as evidenced by the shape of colonies and the assembly of dense actin cytoskeleton networks at their periphery, some level of aggregation was observed. In these aggregates, antibody and phalloidin penetration was restricted and imaging difficult. Therefore, to confirm and quantify pluripotency marker expression over the entire cell population, cell cultures were dissociated and single cells were characterised via FACS.

Flow sorting was carried out, using antibodies against the transcription factors Oct 3/4 and Nanog, as well as the surface markers Stage Specific Embryo Antigen 4 (SSEA-4) and the epitope for TRA-1-60 antibodies as pluripotency markers (Figure 6 and Supplementary Figure S2). In agreement with confocal microscopy images, cell populations displayed high levels of dual marker expression (>95%) for both transcription factors tested, with the rest of the cells mainly expressing Nanog alone. This was the case in all conditions tested, indicating that although the regimen of agitation and size of droplets used had a small impact on cell proliferation and aggregation, it did not impact pluripotency significantly. Further supporting this observation, dual expression of the pluripotency markers SSEA-4 and TRA-1-60R was maintained in all conditions, with >95% cells positive for both markers, whether in dynamic or static conditions. Overall, these results demonstrate that iPSC culture at the surface of microdroplets, mediated by adhesion to strong protein nanosheets, enables their expansion with retention of key pluripotency markers.

**Figure 6.**
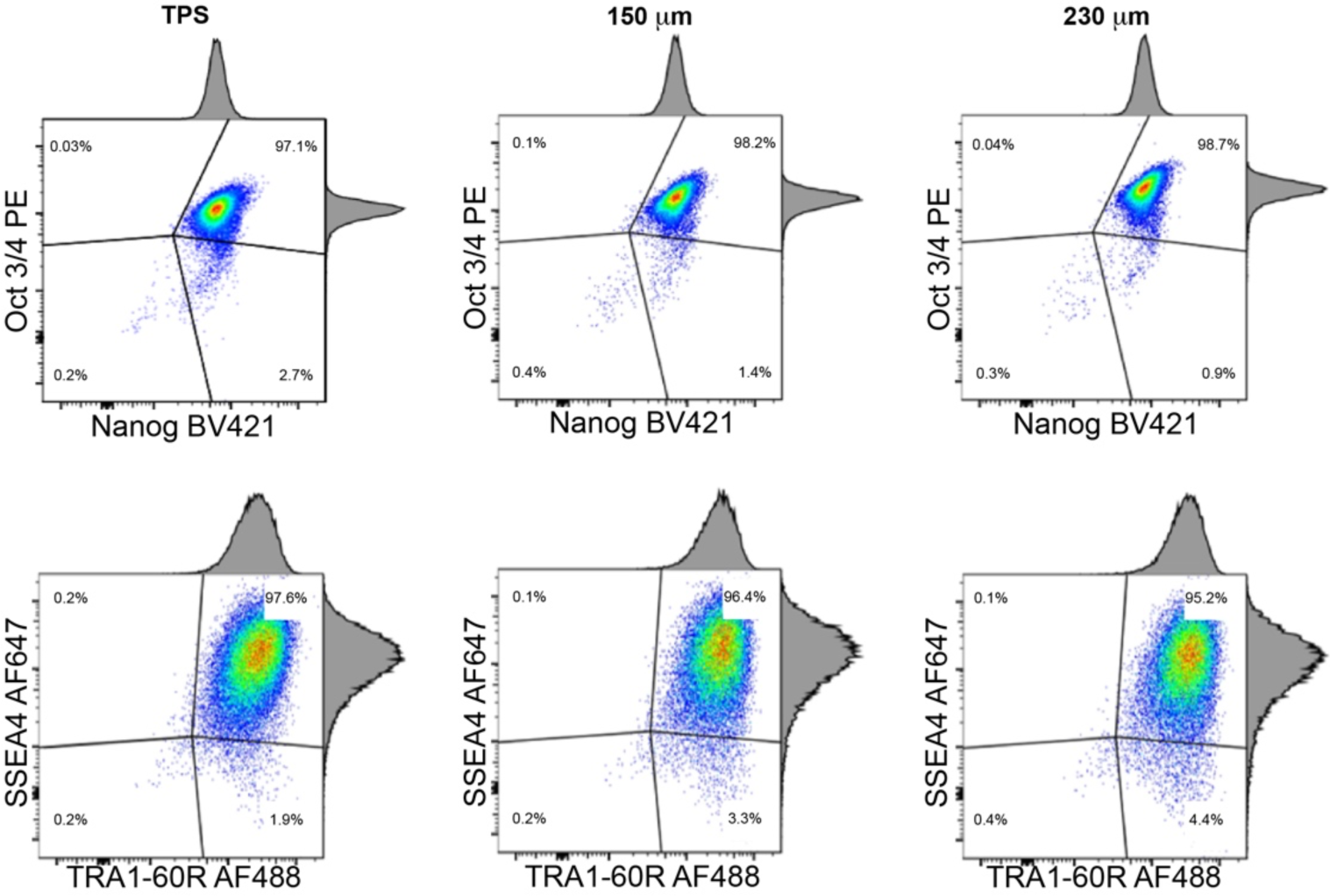
Impact of engineered protein nanosheet-stabilised bioemulsions on iPSC expansion and phenotype characterised by FACS. Flow cytometry of iPSCs grown on βLG-SMCC-stabilised microdroplets (150 and 230 μm; dynamic conditions) for 4 days, quantifying the expression of the pluripotency markers Nanog, Oct 3/4, SSEA-4, and TRA-1-60-R epitope).

### Differentiation of iPSCs on bioemulsions

The ability to induce differentiation of iPSCs towards defined lineages, on bioemulsions, was next examined, taking cardiomyocyte induction as an example. Indeed, iPSC-derived cardiomyocytes have been proposed for cell-based cardiac repair therapies [57,58] and for the development of advanced *in vitro* models [59,60]. A well-established protocol was applied (Figure 1) [43,61], based on initial mesoderm induction using CHIR99021 (2 days), followed by Wnt inhibition using Wnt-C59 (2 days). This was followed by 10 days of culture to allow maturation and glucose starvation for the selection of mature cardiomyocytes. As differentiation on tissue culture plastic is typically associated with detachment and poor efficiency, the differentiation of embryoid bodies was used as reference [62].

Whereas β-catenin was predominantly sequestrated at cell-cell junctions at the end of the expansion period (colocalising with E-cadherin, in particular apically, see Supplementary Figure S3), after priming with CHIR99021, iPSC colonies were found to display clear translocation of β-catenin from cell-cell junctions to the nucleus, already at 24 h (Supplementary Figure S4), and in particular on microdroplets after 48 h of incubation (Supplementary Figure S5). This was also associated with a loss of E-cadherin expression, typical of mesoderm induction [63]. Hence glycogen synthase kinase 3 inhibition resulted in Wnt activation and β -catenin translocation, as expected, including in colonies grown at protein nanosheet-stabilised liquid-liquid interfaces.

After 48 h of mesoderm priming, Wnt was inhibited using the Porcupine O-Acyltransferase (PORCN) inhibitor Wnt-59 [43]. This resulted in the successful restauration of low levels of β-catenin, translocated at junctions, without restauration of E-cadherin expression (Figure 7A).

**Figure 7.**
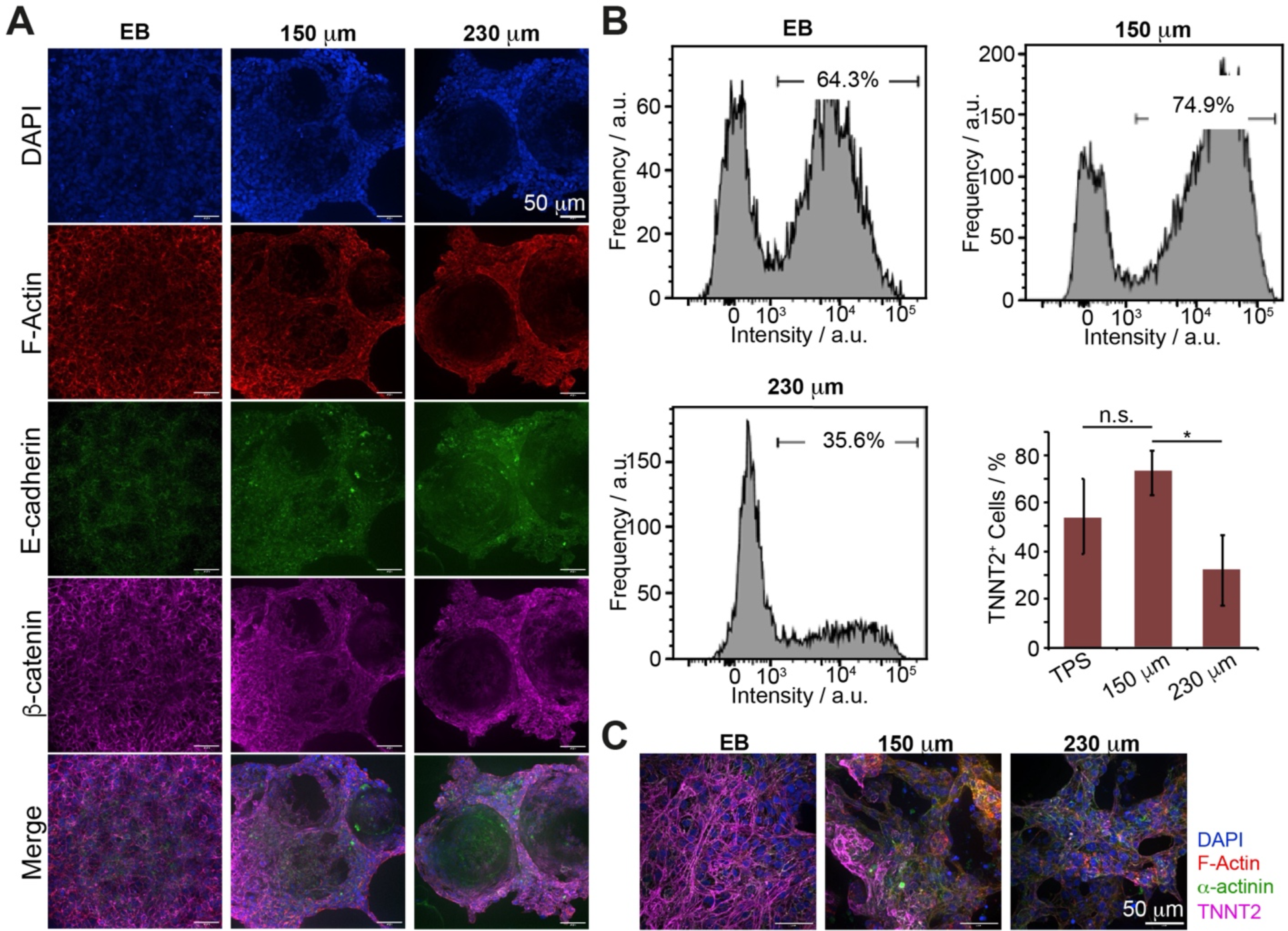
Differentiation of iPSCs into cardiomyocytes on bioemulsions. A. Confocal micrographs of iPSC colonies growing on βLG-SMCC-stabilised microdroplets (diameters of 150 and 230 μm), induced to commit to mesoderm with CHIR99021 (2 days), then Wnt-59 (24 h). Staining for β-catenin (magenta) and E-cadherin (green). Blue, DAPI; Red, Phalloidin/F-actin. B. Flow cytometry (TNNT2) of iPSCs induced to differentiate to cardiomyocytes at corresponding bioemulsions for 10 days, followed by metabolic selection for 6 days. Corresponding quantification of positive cell populations. n = 4. n.s., not significant; *, p < 0.05. C. Confocal micrographs of resulting cultures. Staining for TNNT2 (magenta) and α-actinin (green). Blue, DAPI; Red, Phalloidin/F-actin.

Following this second inhibition period, primed cells were cultured in differentiation medium 3 (CDM3; Figure 1) for 10 days, prior to glucose starvation for metabolic selection of mature cardiomyocytes. The resulting cell cultures were immunostained and imaged at days 7 and 10 to evaluate the expression of the cardiac lineage markers TNNT2 and α-actinin (Supplementary Figure S6) [43,64]. At day 7, TNNT2 expression was found to have initiated relatively extensively on cultures at βLG-SMCC-stabilised liquid-liquid interfaces, whereas this remained at a budding stage on tissue culture plastic. However, by day 10, culture on solid substrates had caught up. To evaluate TNNT2 expression following metabolic selection, TNNT2 expression was monitored by flow cytometry (Figure 7B). Culture at the surface of large microdroplets (150 μm) was found to lead to comparable populations of TNNT2 positive cells to culture on tissue culture plastic (TPS) and significantly higher levels of expression compared to cultures on 230 μm droplets. This was further confirmed by immunostaining (Figure 7C).

Finally, to confirm the maturity of cardiomyocytes generated at liquid-liquid interfaces stabilised by βLG/SMCC nanosheets, time-lapse imaging of colonies was carried out (Figure S7A). Beating colonies were clearly observed throughout the culture. Substrate deformation was even apparent, as colonies exerted forces that resulted in the wrinkling of nanosheets (Supplementary Figure S7B). Hence nanosheet-stabilised liquid-liquid interfaces of microdroplets sustained the differentiation of iPSCs towards functional cardiomyocytes.

## Conclusions

The ability of cells to sense and respond to mechanical and other physical properties of their environment has been shown to impact on a broad range of processes and regulate the differentiation of multipotent as well as pluripotent stem cells [65,66]. The ability to maintain iPSC expansion capacity and pluripotency markers expression at liquid-liquid interfaces stabilised by protein nanosheets further demonstrates the impact that nanoscale mechanical properties of the cell microenvironment plays in maintaining stem cell phenotype. Hence, although low viscosity oil substrates were used for the culture of iPSCs in this study, the strong elastic interfacial shear mechanics resulting from protein nanosheet assembly and crosslinking enabled resistance to cell-mediated forces exerted during colony development.

In turn, the compatibility of such assemblies with microdroplet microfluidic platforms enabled the generation of bioemulsions with well-defined microdroplet sizes, to investigate the impact that such parameter has on the expansion and differentiation of iPSCs. Whereas little impact was observed on the retention of pluripotency markers, cardiomyocyte differentiation was more significantly impacted. Beyond interfacial shear mechanical properties, the curvature of droplets and the tendency of multiple colonies to aggregates is the likely cause for such observation. Indeed, substrate curvature was found to impact cytoskeleton assembly, cell motility and the differentiation of mesenchymal stem cells [67,68]. Although TNNT2 expression was not completely supressed by culture on large microdroplets, it could be proposed that fine changes in the balance of cell-matrix adhesion, contractility and cell-cell junction maturation (and associated β-catenin sequestration) could underpin such differential regulation of iPSC differentiation. Similar crosstalks between cell-cell and cell-matrix adhesions have been reported to regulate processes such as differentiation and motility, at the surface of compliant hydrogels [63,69].

Culture on liquid substrate is particularly attractive to tackle some of the challenges facing cell delivery for tissue engineering and regenerative medicine. Indeed, unlike culture on solid substrates, including plastic microcarriers, that typically requires cell processing and detachment, and may result in contamination of cell products with microplastics, bioemulsions could allow the direct implantation of adherent cell colonies. Although fluorinated oils are unlikely to be translatable for implantation, they present a proof-of-concept of the ability to regulate stem cell phenotype and differentiation at liquid-liquid interfaces, with suitable protein nanosheet engineering. In addition, a recent report demonstrated the capacity to maintain mesenchymal stem cell expansion at the surface of a range of non-fluorinated liquids, including silicone, mineral and plant-based oils [70]. Therefore bioemulsions present unique opportunities to transform cell manufacturing and cell delivery processes and protocols.

## Acknowledgement

We thank the European Research Council (ProLiCell, 772462) for support.

## Reference

[1] Gurdon, J.B. (1962). The developmental capacity of nuclei taken from intestinal epithelium cells of feeding tadpoles. Journal of embryology and experimental morphology. DOI: 10.1242/dev.10.4.622.

[2] Gurdon, J.B. et al. (1958). Sexually mature individuals of Xenopus laevis from the transplantation of single somatic nuclei. Nature. DOI: 10.1038/182064a0.

[3] Takahashi, K. and Yamanaka, S. (2006). Induction of Pluripotent Stem Cells from Mouse Embryonic and Adult Fibroblast Cultures by Defined Factors. Cell. DOI: 10.1016/j.cell.2006.07.024.

[4] Takahashi, K. and Yamanaka, S. (2016). A decade of transcription factor-mediated reprogramming to pluripotency. Nature Reviews Molecular Cell Biology. DOI: 10.1038/nrm.2016.8.

[5] Junying, Y. et al. (2009). Human induced pluripotent stem cells free of vector and transgene sequences. Science. DOI: 10.1126/science.1172482.

[6] Takahashi, K. et al. (2007). Induction of Pluripotent Stem Cells from Adult Human Fibroblasts by Defined Factors. Cell. DOI: 10.1016/j.cell.2007.11.019.

[7] Yu, J. et al. (2007). Induced pluripotent stem cell lines derived from human somatic cells. Science. DOI: 10.1126/science.1151526.

[8] Chen, G. et al. (2011). Chemically defined conditions for human iPSC derivation and culture. Nature Methods. DOI: 10.1038/nmeth.1593.

[9] Amabile, G. and Meissner, A. (2009). Induced pluripotent stem cells: current progress and potential for regenerative medicine. Trends in Molecular Medicine. DOI: 10.1016/j.molmed.2008.12.003.

[10] Martin, U. (2017). Therapeutic application of pluripotent stem cells: Challenges and risks. Frontiers in Medicine. DOI: 10.3389/fmed.2017.00229.

[11] Reardon, S. and Cyranoski, D. (2014). Japan stem-cell trial stirs envy. Nature. DOI: 10.1038/513287a.

[12] Mandai, M. et al. (2017). Autologous Induced Stem-Cell–Derived Retinal Cells for Macular Degeneration. New England Journal of Medicine. DOI: 10.1056/nejmoa1608368.

[13] Lancaster, M.A. et al. (2013). Cerebral organoids model human brain development and microcephaly. Nature. DOI: 10.1038/nature12517.

[14] Birey, F. et al. (2017). Assembly of functionally integrated human forebrain spheroids. Nature. DOI: 10.1038/nature22330.

[15] Mariani, J. et al. (2015). FOXG1-Dependent Dysregulation of GABA/Glutamate Neuron Differentiation in Autism Spectrum Disorders. Cell. DOI: 10.1016/j.cell.2015.06.034.

[16] Qian, X. et al. (2016). Brain-Region-Specific Organoids Using Mini-bioreactors for Modeling ZIKV Exposure. Cell. DOI: 10.1016/j.cell.2016.04.032.

[17] Takebe, T. et al. (2013). Vascularized and functional human liver from an iPSC-derived organ bud transplant. Nature. DOI: 10.1038/nature12271.

[18] Gao, D. et al. (2014). Organoid cultures derived from patients with advanced prostate cancer. Cell. DOI: 10.1016/j.cell.2014.08.016.

[19] Spence, J.R. et al. (2011). Directed differentiation of human pluripotent stem cells into intestinal tissue in vitro. Nature. DOI: 10.1038/nature09691.

[20] Negoro, R. et al. (2018). Efficient Generation of Small Intestinal Epithelial-like Cells from Human iPSCs for Drug Absorption and Metabolism Studies. Stem Cell Reports. DOI: 10.1016/j.stemcr.2018.10.019.

[21] Dekkers, J.F. et al. (2016). Characterizing responses to CFTR-modulating drugs using rectal organoids derived from subjects with cystic fibrosis. Science Translational Medicine. DOI: 10.1126/scitranslmed.aad8278.

[22] Kim, J. et al. (2013). An iPSC Line from Human Pancreatic Ductal Adenocarcinoma Undergoes Early to Invasive Stages of Pancreatic Cancer Progression. Cell Reports. DOI: 10.1016/j.celrep.2013.05.036.

[23] Ramme, A.P. et al. (2018). Towards an autologous iPSC-derived patient-on-a-chip. bioRxiv.

[24] Huch, M. et al. (2017). The hope and the hype of organoid research. Development (Cambridge). DOI: 10.1242/dev.150201.

[25] Borys, B.S. et al. (2020). Optimized serial expansion of human induced pluripotent stem cells using low-density inoculation to generate clinically relevant quantities in vertical-wheel bioreactors. Stem Cells Translational Medicine. DOI: 10.1002/sctm.19-0406.

[26] Badenes, S.M. et al. (2016). Defined essential 8′ medium and vitronectin efficiently support scalable xeno-free expansion of human induced pluripotent stem cells in stirred microcarrier culture systems. PLoS ONE. DOI: 10.1371/journal.pone.0151264.

[27] Van Wezel, A.L. (1967). Growth of cell-strains and primary cells on micro-carriers in homogeneous culture [17]. Nature. DOI: 10.1038/216064a0.

[28] Tavassoli, H. et al. (2018). Large-scale production of stem cells utilizing microcarriers: A biomaterials engineering perspective from academic research to commercialized products. Biomaterials. DOI: 10.1016/j.biomaterials.2018.07.016.

[29] Sart, S. et al. (2013). Engineering stem cell fate with biochemical and biomechanical properties of microcarriers. Biotechnology Progress. DOI: 10.1002/btpr.1825.

[30] Zhang, J. et al. (2015). Thermo-responsive microcarriers based on poly(N-isopropylacrylamide). European Polymer Journal. DOI: 10.1016/j.eurpolymj.2015.04.013.

[31] Tang, Z. et al. (2012). Temperature-responsive polymer modified surface for cell sheet engineering. Polymers. DOI: 10.3390/polym4031478.

[32] Kong, D. et al. (2022). Impact of the multiscale viscoelasticity of quasi-2D self-assembled protein networks on stem cell expansion at liquid interfaces. Biomaterials. DOI: 10.1016/j.biomaterials.2022.121494.

[33] Chrysanthou, A. et al. (2022). Supercharged Protein Nanosheets for Cell Expansion on Bioemulsions. ACS Applied Materials and Interfaces. DOI: 10.1021/acsami.2c20188.

[34] Peng, L. et al. (2023). Mesenchymal Stem Cells Sense the Toughness of Nanomaterials and Interfaces. Advanced Healthcare Materials. DOI: 10.1002/adhm.202203297.

[35] Jia, X. et al. (2022). Adaptive liquid interfaces induce neuronal differentiation of mesenchymal stem cells through lipid raft assembly. Nature Communications. DOI: 10.1038/s41467-022-30622-y.

[36] Jia, X. et al. (2020). Adaptive Liquid Interfacially Assembled Protein Nanosheets for Guiding Mesenchymal Stem Cell Fate. Advanced Materials. DOI: 10.1002/adma.201905942.

[37] Kong, D. et al. (2017). The culture of HaCaT cells on liquid substrates is mediated by a mechanically strong liquid-liquid interface. Faraday Discussions. DOI: 10.1039/c7fd00091j.

[38] Kong, D. et al. (2018). Protein Nanosheet Mechanics Controls Cell Adhesion and Expansion on Low-Viscosity Liquids. Nano Letters. DOI: 10.1021/acs.nanolett.7b05339.

[39] Chrysanthou, A. et al. (2022). Co-Surfactant-Free Bioactive Protein Nanosheets for the Stabilization of Bioemulsions Enabling Adherent Cell Expansion. Biomacromolecules. DOI: 10.1021/acs.biomac.2c01289.

[40] Kong, D. et al. (2018). Protein Nanosheet Mechanics Controls Cell Adhesion and Expansion on Low-Viscosity Liquids. Nano Letters. DOI: 10.1021/acs.nanolett.7b05339.

[41] Peng, L. and Gautrot, J.E. (2021). Long term expansion profile of mesenchymal stromal cells at protein nanosheet-stabilised bioemulsions for next generation cell culture microcarriers. Materials Today Bio. DOI: 10.1016/j.mtbio.2021.100159.

[42] Kong, D. et al. (2018). Stem Cell Expansion and Fate Decision on Liquid Substrates Are Regulated by Self-Assembled Nanosheets. ACS Nano. DOI: 10.1021/acsnano.8b03865.

[43] Burridge, P.W. et al. (2014). Chemically defined generation of human cardiomyocytes-supplementary information. Nature Methods.

[44] Tamkun, J.W. and Hynes, R.O. (1983). Plasma fibronectin is synthesized and secreted by hepatocytes. Journal of Biological Chemistry. DOI: 10.1016/s0021-9258(18)32672-3.

[45] Preissner, K.T. (1991). Structure and biological role of vitronectin. Annual Review of Cell Biology. DOI: 10.1146/annurev.cb.07.110191.001423.

[46] Fuller, G.G. and Vermant, J. (2012). Complex Fluid-Fluid Interfaces: Rheology and Structure. Annual Review of Chemical and Biomolecular Engineering. DOI: 10.1146/annurev-chembioeng-061010-114202.

[47] Vandebril, S. et al. (2010). A double wall-ring geometry for interfacial shear rheometry. Rheologica Acta. DOI: 10.1007/s00397-009-0407-3.

[48] Megone, W. et al. (2021). Extreme reversal in mechanical anisotropy in liquid-liquid interfaces reinforced with self-assembled protein nanosheets. Journal of Colloid and Interface Science. DOI: 10.1016/j.jcis.2021.03.055.

[49] Yu, Z. et al. (2018). Droplet-based microfluidic analysis and screening of single plant cells. PLoS ONE. DOI: 10.1371/journal.pone.0196810.

[50] Baret, J.C. (2012). Surfactants in droplet-based microfluidics. Lab on a Chip. DOI: 10.1039/c1lc20582j.

[51] Garstecki, P. et al. (2005). Mechanism for flow-rate controlled breakup in confined geometries: A route to monodisperse emulsions. Physical Review Letters. DOI: 10.1103/PhysRevLett.94.164501.

[52] Anna, S.L. et al. (2003). Formation of dispersions using “flow focusing” in microchannels. Applied Physics Letters. DOI: 10.1063/1.1537519.

[53] Nunes, J.K. et al. (2013). Dripping and jetting in microfluidic multiphase flows applied to particle and fibre synthesis. Journal of Physics D: Applied Physics. DOI: 10.1088/0022-3727/46/11/114002.

[54] Rodrigues, A.L. et al. (2019). Dissolvable Microcarriers Allow Scalable Expansion And Harvesting Of Human Induced Pluripotent Stem Cells Under Xeno-Free Conditions. Biotechnology Journal. DOI: 10.1002/biot.201800461.

[55] Petry, F. et al. (2016). Manufacturing of Human Umbilical Cord Mesenchymal Stromal Cells on Microcarriers in a Dynamic System for Clinical Use. Stem Cells International. DOI: 10.1155/2016/4834616.

[56] Badenes, S.M. et al. (2016). Microcarrier-based platforms for in vitro expansion and differentiation of human pluripotent stem cells in bioreactor culture systems. Journal of Biotechnology. DOI: 10.1016/j.jbiotec.2016.07.023.

[57] Rojas, S. V. et al. (2017). Transplantation of purified iPSC-derived cardiomyocytes in myocardial infarction. PLoS ONE. DOI: 10.1371/journal.pone.0173222.

[58] Tu, C. and Zoldan, J. (2018). Moving iPSC-Derived Cardiomyocytes Forward to Treat Myocardial Infarction. Cell Stem Cell. DOI: 10.1016/j.stem.2018.08.011.

[59] Sinnecker, D. et al. (2013). Induced pluripotent stem cell-derived cardiomyocytes: A versatile tool for arrhythmia research. Circulation Research. DOI: 10.1161/CIRCRESAHA.112.268623.

[60] Di Cio, S. et al. Vascularised Cardiac Spheroids-on-a-Chip for Testing the Toxicity of Therapeutics. BioRxiv.

[61] Ueno, S. et al. (2007). Biphasic role for Wnt/β-catenin signaling in cardiac specification in zebrafish and embryonic stem cells. Proceedings of the National Academy of Sciences of the United States of America. DOI: 10.1073/pnas.0702859104.

[62] Azarin, S.M. et al. (2012). Modulation of Wnt/β-catenin signaling in human embryonic stem cells using a 3-D microwell array. Biomaterials. DOI: 10.1016/j.biomaterials.2011.11.070.

[63] Przybyla, L. et al. (2016). Tissue Mechanics Orchestrate Wnt-Dependent Human Embryonic Stem Cell Differentiation. Cell Stem Cell. DOI: 10.1016/j.stem.2016.06.018.

[64] Palpant, N.J. et al. (2017). Generating high-purity cardiac and endothelial derivatives from patterned mesoderm using human pluripotent stem cells. Nature Protocols. DOI: 10.1038/nprot.2016.153.

[65] Guilak, F. et al. (2009). Control of Stem Cell Fate by Physical Interactions with the Extracellular Matrix. Cell Stem Cell. DOI: 10.1016/j.stem.2009.06.016.

[66] Di Cio, S. and Gautrot, J.E. (2016). Cell sensing of physical properties at the nanoscale: Mechanisms and control of cell adhesion and phenotype. Acta Biomaterialia. DOI: 10.1016/j.actbio.2015.11.027.

[67] Pieuchot, L. et al. (2018). Curvotaxis directs cell migration through cell-scale curvature landscapes. Nature Communications. DOI: 10.1038/s41467-018-06494-6.

[68] Callens, S.J.P. et al. (2020). Substrate curvature as a cue to guide spatiotemporal cell and tissue organization. Biomaterials. DOI: 10.1016/j.biomaterials.2019.119739.

[69] Borghi, N. et al. (2010). Regulation of cell motile behavior by crosstalk between cadherin- and integrin-mediated adhesions. Proceedings of the National Academy of Sciences of the United States of America. DOI: 10.1073/pnas.1002662107.

[70] Peng, L. et al. (2023). Growth of mesenchymal stem cells at the surface of silicone, mineral and plant-based oils. Biomedical Materials. DOI: 10.1088/1748-605x/acbdda.

