## Supplementary Data for "Strong Elastic Protein Nanosheets Enable the Culture and Differentiation of Induced Pluripotent Stem Cells on Microdroplets"

### Supplementary Information

Elijah Mojares[1], Alexandra Chrysanthou[1] and Julien E. Gautrot[1*]

[1] School of Engineering and Materials Science, Queen Mary University of London, Mile End Road, London E1 4NS, United Kingdom.

* Correspondence:

Julien E. Gautrot

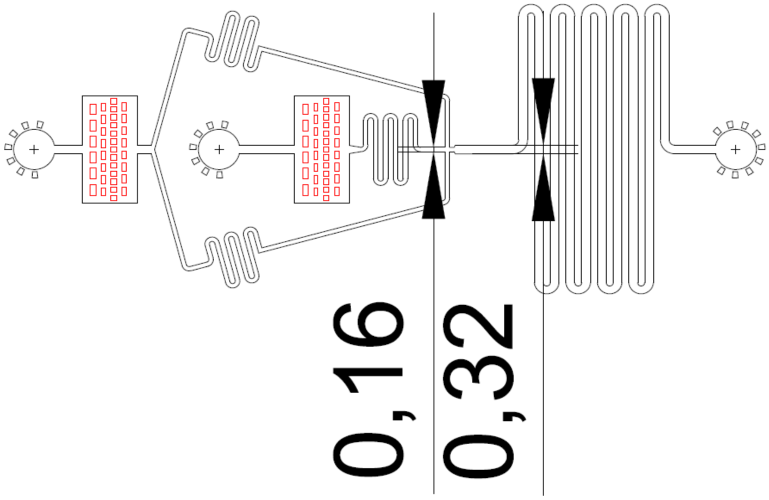


**Supplementary Figure S1.** Schematic of the microdroplet microfluidic chip used to produced nanosheet-stabilised bioemulsions.

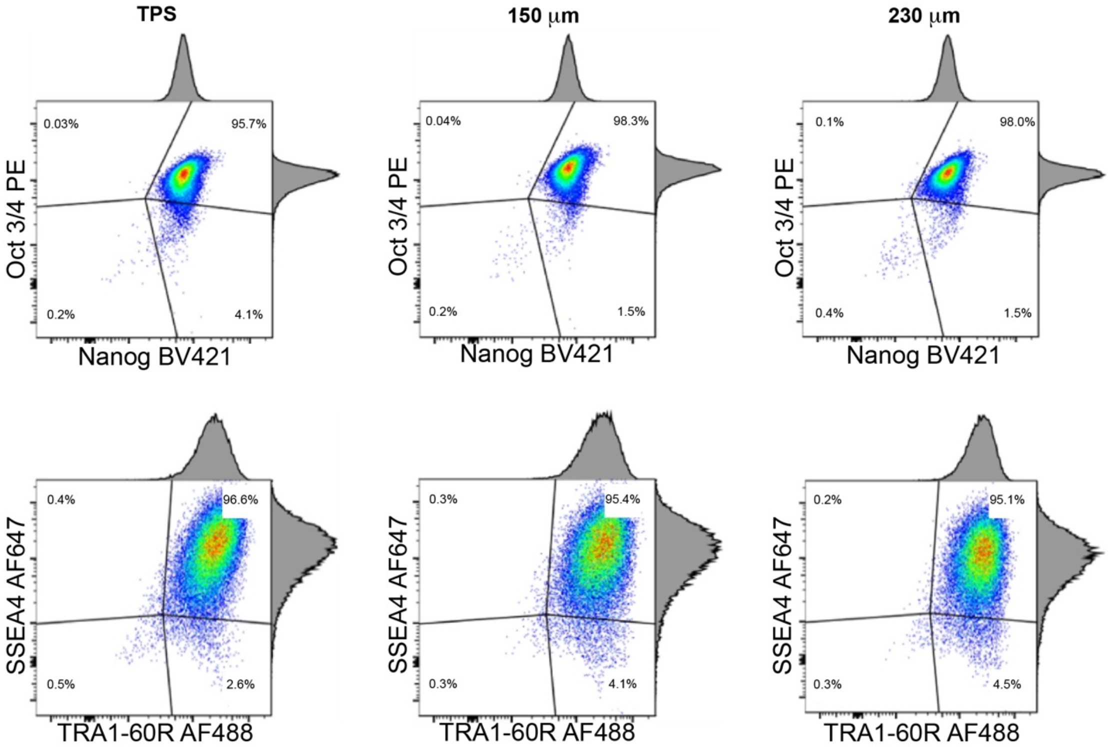


**Supplementary Figure S2.** Impact of engineered protein nanosheet-stabilised bioemulsions on iPSC expansion and phenotype characterised by FACS. Flow cytometry of iPSCs grown on βLG-SMCC-stabilised microdroplets (150 and 230 μm; static conditions) for 4 days, quantifying the expression of the pluripotency markers Nanog, Oct 3/4, SSEA-4, and TRA-1-60-R epitope).

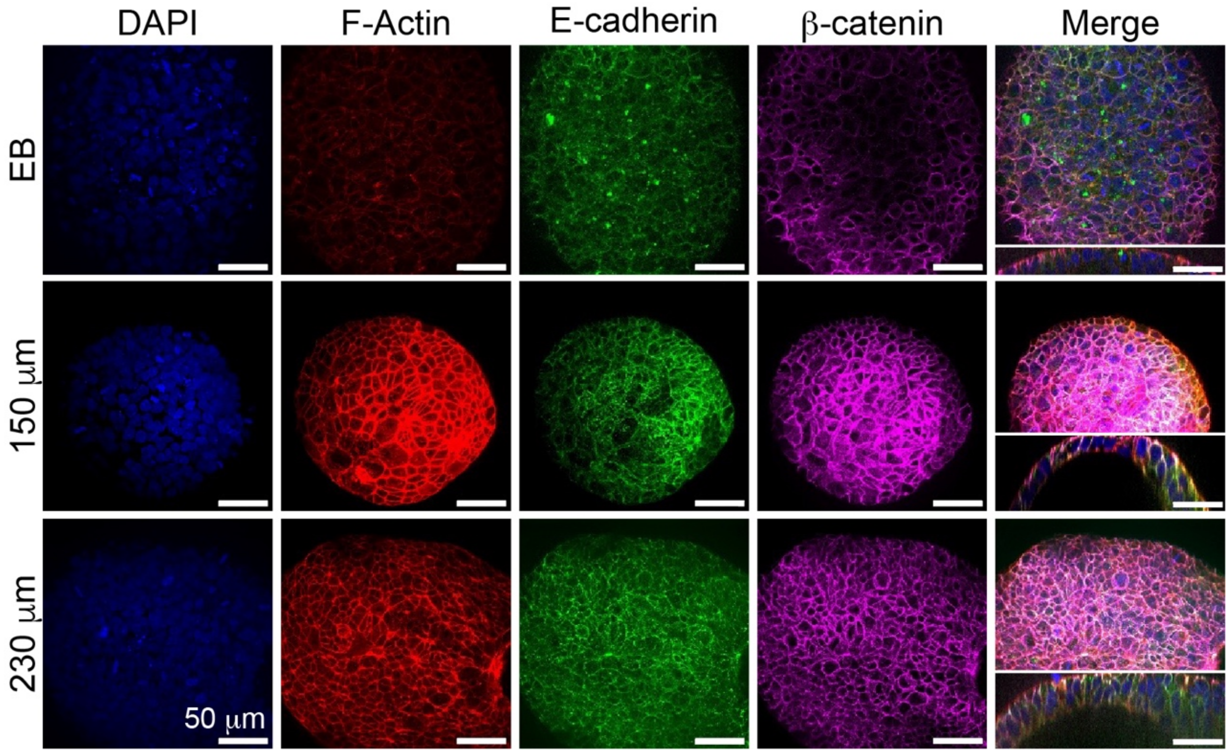


**Supplementary Figure S3.** Confocal micrographs of iPSC colonies growing on βLG-SMCC-stabilised microdroplets (diameters of 150 and 230 μm), in dynamic conditions, compared to embryoid bodies (EB). Staining for β-catenin (magenta) and E-cadherin (green). Blue, DAPI; Red, Phalloidin/F-actin.

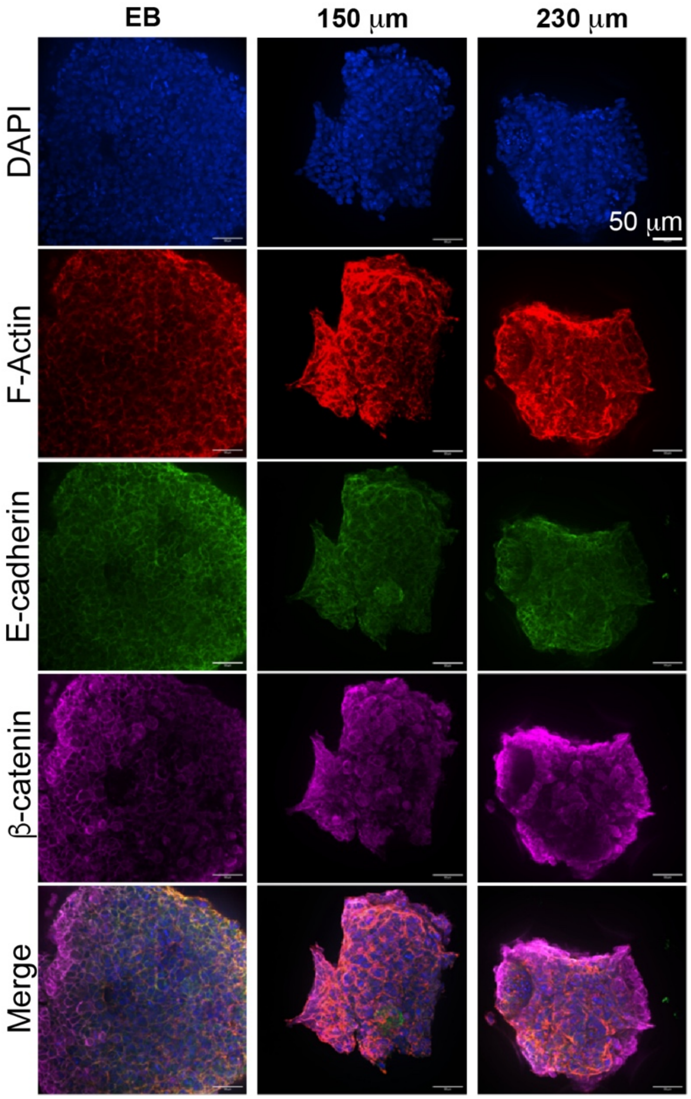


**Supplementary Figure S4.** Confocal micrographs of iPSC colonies growing on βLG-SMCC-stabilised microdroplets (diameters of 150 and 230 μm), compared to embryoid bodies (EB), after 24 h of priming with CHIR99021 (6 μM). Staining for β-catenin (magenta) and E-cadherin (green). Blue, DAPI; Red, Phalloidin/F-actin.

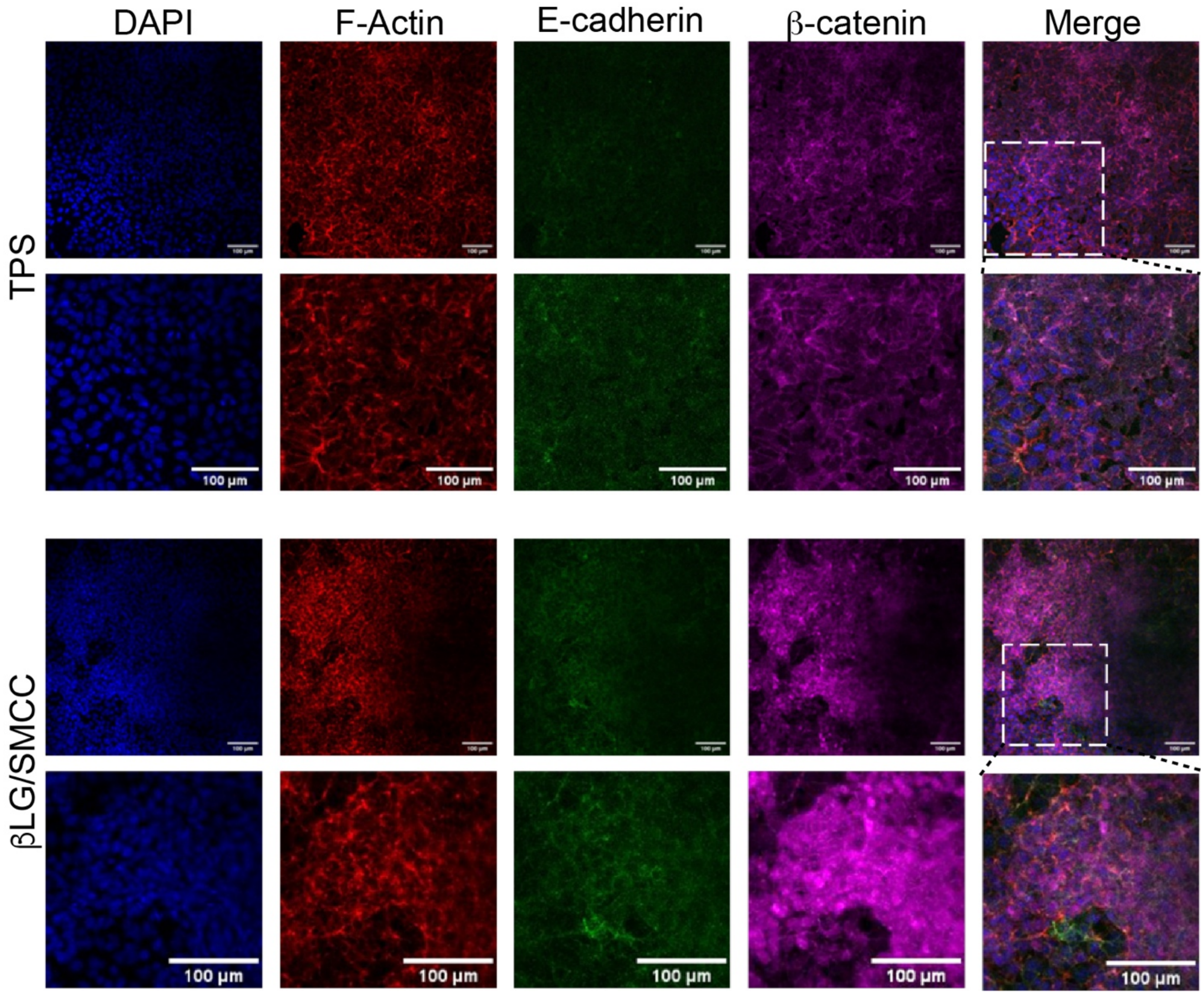


**Supplementary Figure S5.** Confocal micrographs of iPSC colonies growing on βLG-SMCC-stabilised microdroplets (diameters of 150 and 230 μm), compared to embryoid bodies (EB), after 48 h of priming with CHIR99021 (6 μM). Staining for β-catenin (magenta) and E-cadherin (green). Blue, DAPI; Red, Phalloidin/F-actin.

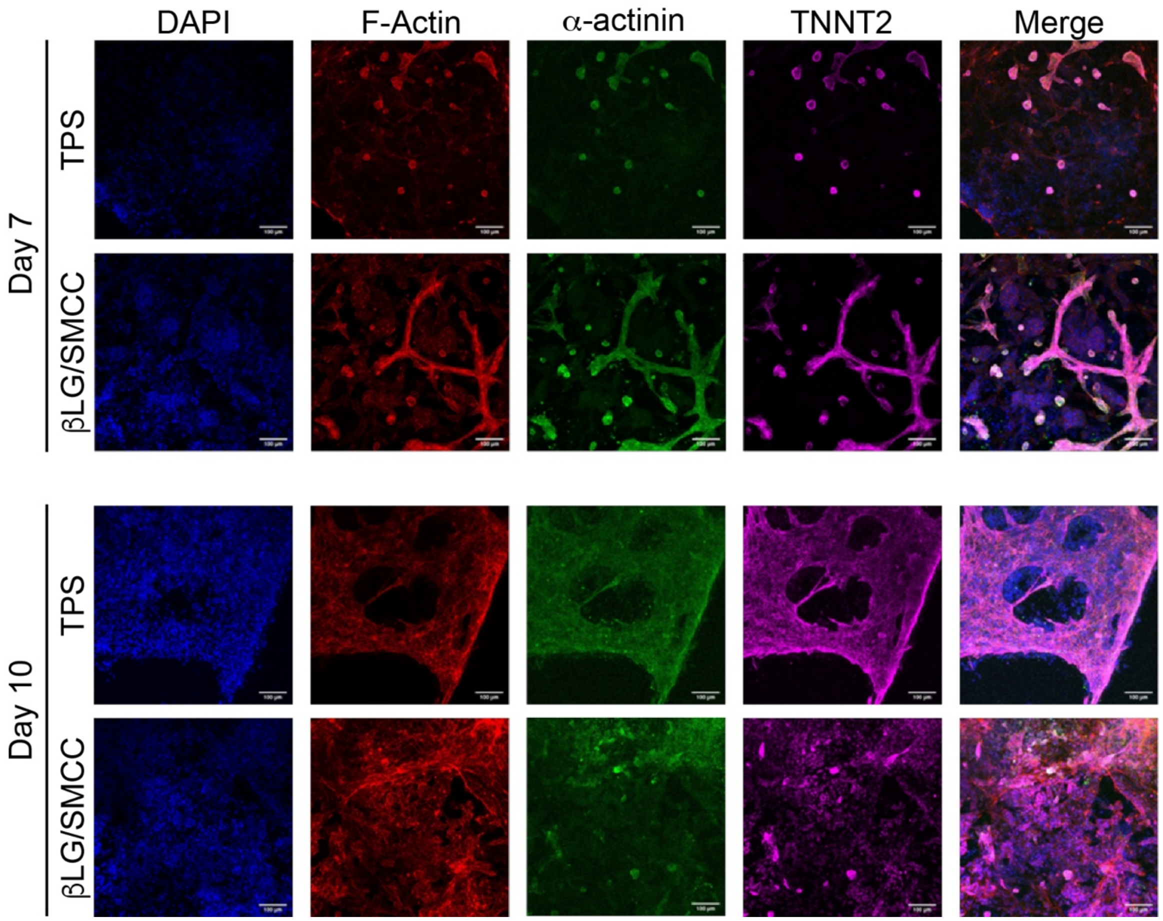


**Supplementary Figure S6.** Confocal micrographs of iPSC colonies growing on βLG-SMCC-stabilised Novec 7500-PBS interfaces and TPS, after priming and 7 and 10 days of differentiation in CDM3. Staining for α-actinin (magenta) and TNNT2 (green). Blue, DAPI; Red, Phalloidin/F-actin.

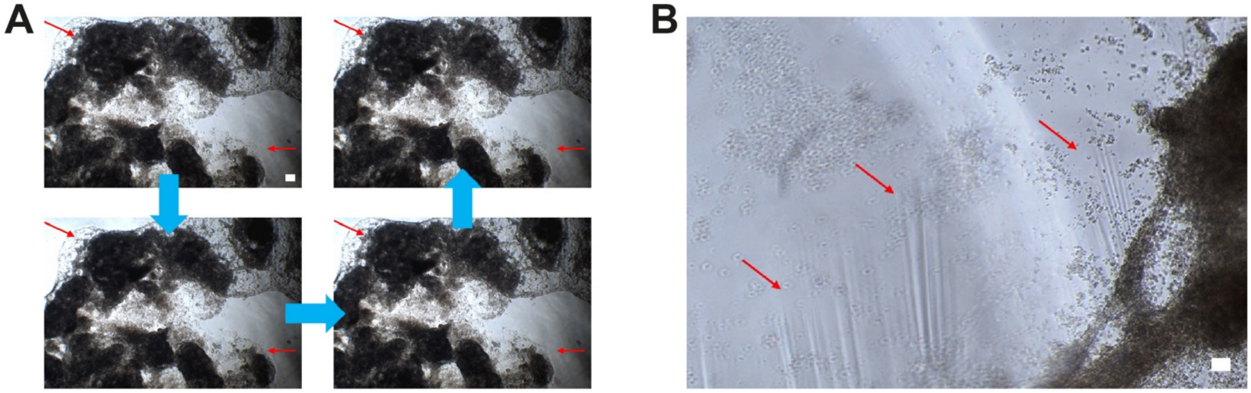


**Supplementary Figure S6.** Beating cardiomyocytes generated on a βLG-SMCC nanosheet stabilised liquid-liquid interface.

**Supplementary Video S1.** Beating cardiomyocytes generated on a βLG-SMCC-stabilised mirodroplets.
